# CD4^+^ and CD8^+^ regulatory T cells characterization in the rat using a unique transgenic *Foxp3-EGFP* model

**DOI:** 10.1101/2021.12.09.471889

**Authors:** Séverine Ménoret, Laurent Tesson, Séverine Remy, Victor Gourain, Céline Sérazin, Claire Usal, Aude Guiffes, Vanessa Chenouard, Laure-Hélène Ouisse, Malika Gantier, Jean-Marie Heslan, Cynthia Fourgeux, Jeremie Poschmann, Carole Guillonneau, Ignacio Anegon

## Abstract

**Background:** CD4^+^ and CD8^+^ regulatory T cells (Treg) in diverse species include different subsets from different origins. In all species, CD8^+^ Treg have been poorly characterized. CD4^+^ and CD8^+^ Treg in rats have only partially been characterized and there is no rat model in which FOXP3^+^ Treg are genetically tagged.

**Results:** We generated a rat transgenic line using the CRISPR/Cas9 system in which EGFP was inserted in frame on the 3’ end of the *Foxp3* gene using a 2A self-cleaving peptide. EGFP was exclusively expressed by CD4^+^ and CD8^+^ T cells in similar proportion as observed with anti-FOXP3 antibodies. CD4^+^EGFP^+^ Treg were 5-10 times more frequent than CD8^+^EGFP^+^ Treg. CD4^+^ and CD8^+^ EGFP^+^ Treg expressed both the CD25^high^CD127^low^CD45RC^low/-^ markers. The suppressive activity of CD4^+^ and CD8^+^ Treg was largely confined to EGFP^+^ cells. RNAseq analyses showed similarities but also differences among CD4^+^ and CD8^+^ EGFP^+^ cells and provided the first description of the natural FOXP3^+^ CD8^+^ Treg transcriptome. In vitro culture of CD4^+^ and CD8^+^ EGFP^-^ cells with TGFbeta and IL-2 resulted in the induction of EGFP^+^ Treg. Preferential expansion of CD4^+^ and CD8^+^ EGFP^+^ Treg could be detected upon in vivo administration of a low dose of IL-2.

**Conclusions:** This new and unique *Foxp3-EGFP* rat line constitutes a useful model to identify and isolate viable natural and induced CD4^+^ and CD8^+^ Treg. Additionally, it allows to identify new molecules expressed in CD8^+^ Treg that may allow to better define their phenotype and function not only in rats but also in other species.

## Introduction

Regulatory T cells (Treg) play a central role in controlling immune effector mechanisms (1–3). Both, CD4^+^ and CD8^+^ Treg have been repeatedly described in humans, non-human primates and rodents (3–5). CD4^+^FOXP3^+^ Treg can be natural (nTregs), i.e. physiologically developed in the thymus and then migrate to the periphery (3). The existence of natural CD8^+^FOXP3^+^ Treg has not been yet formally demonstrated (4). At the same time, CD4^+^ and CD8^+^FOXP3^+^ Treg can be induced from FOXP3^-^ T (iTreg) in pathophysiological situations or following treatments, either *in vivo* or *in vitro* (1, 3, 4, 6, 7). Additionally, there are also CD4^+^ and CD8^+^ iTreg that are FOXP3^-^ (1, 4, 8). The FOXP3 transcription factor is essential for the function of canonical natural CD4^+^ Treg (1). CD8^+^ Treg is and heterogeneous cell population, as CD4^+^ Treg, but the phenotype and function of natural CD8^+^FOXP3^+^ Treg is less well defined than for CD4^+^FOXP3^+^ Treg (4). We and others have described in rats and humans a population of cells CD8^+^ Treg that is defined by the low or absent expression of CD45RC (CD8^+^CD45RC^low/-^)(7, 9–12). In mice, although CD45RC has been used to purify CD4^+^ and CD8^+^ (13) Treg little characterization of these cells has been done. In rats and humans all CD4^+^ and CD8^+^ Treg suppressive activity is comprised among the CD45RC^low/-^ fraction but the FOXP3^+^ cells are only a small fraction of this population. Although CD4^+^ Treg in all species are comprised in the CD25^high^CD127^low^ fraction of CD45RC^low/-^ cells, for CD8^+^ Treg there is no consensus on the markers useful to further refine the analysis of the CD45RC^low/-^ population and to analyze if the suppressive activity is exclusively within the FOXP3^+^ fraction (4, 7).

Mouse FOXP3^+^CD4^+^ Treg have been extensively analyzed using transgenic animals in which the *Foxp3* promoter controls reporter genes (14–20) allowing the sorting of viable CD4^+^ *Foxp3-GFP^+^* Treg. CD8^+^FOXP3^+^ cells were described but an in dept analysis of their phenotype and function was not reported (15–18).

Rat CD4^+^ and CD8^+^ Treg are in general less well defined than human and mouse Treg and comparison of Treg among different species are very rare and only on the T CD4^+^ lineage (5). There is no description of rat transgenic lines that allow to define FOXP3^+^ cell lineages.

The immune system of rats have several characteristics that make them more similar to humans than mice, such as expression of MHC-II molecules by T cells, CD4 expression by a subset of DCs and normal levels of complement (21, 22). Additionally, certain human immune diseases are better modelized in rat than in mouse models with the same genetic mutations (22), such as for ankylosing spondylopathy (23) and autoimmune polyendocrinopathy candidiasis ectodermal dystrophy (APECED) (24), in *HLA-B27* transgenic and in *Aire* knockout rats, respectively.

Thus, there is need to better define Treg in the rat, specially the CD8^+^ Treg population and to dispose of animals with Foxp3-tagged Treg that can be used to study Treg in different situations.

Using CRISPR/Cas9 and transgenic technologies we generated a rat line with a knockin of EGFP in the 3’ of the *Foxp3* gene allowing the natural expression of FOXP3 from the targeted allele with the expression of EGFP under the control of the endogenous *Foxp3* promoter. The phenotype, suppressive function and transcriptomic of both CD4^+^ and CD8^+^ EGFP^+^ Treg were analyzed. The transcriptomic analysis of CD8^+^EGFP^+^ Treg is to the best of our knowledge the first one to be described. Using EGFP as a marker we show expansion of both CD4^+^ and CD8^+^ Treg following IL-2 treatment in vivo and generation of iTreg in vitro.

## Results

### Generation of transgenic rats with EGFP knockin into the *Foxp3* gene

To mark FOXP3-expressing cells we inserted the EGFP coding sequence in-frame and before the stop codon of the *Foxp3* gene (**Fig. 1A**). EGFP was preceded by a T2A sequence that allowed self-splicing of a monocistronic *Foxp3-EGFP* mRNA. Thus *Foxp3*-expressing cells will show simultaneous and independent expression of FOXP3 and EGFP under the control of the endogenous *Foxp3* promoter (referred to as *Foxp3-EGFP* rats below).

**Figure 1.**
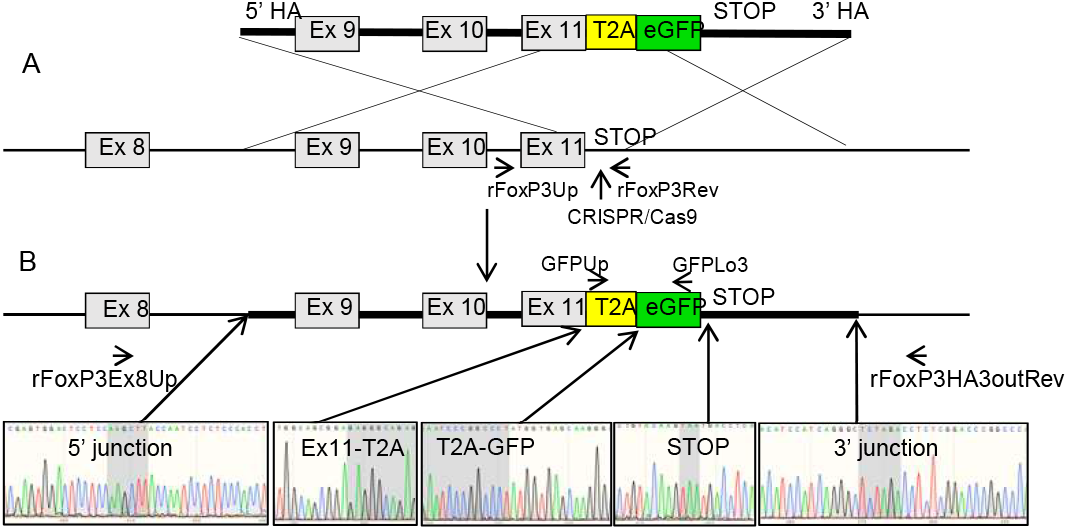
Schematic representation of the GFP KI strategy and genotyping of potential *Foxp3-EGFP* founders. **A.** (upper diagram) A double stranded DNA donor was generated containing a EGFP sequence preceded by a self-splicing T2A sequence both flanked in 5’ and 3’ homology arms for the indicated *Foxp3* sequences. (lower diagram). Upon microinjection of a sgRNA designed to recognize a DNA sequence immediately 3’ of the stop codon, Cas9 protein and donor DNA, cleavage of the *Foxp3* gene favors homologous-recombination of the donor DNA sequences. **B.** (upper diagram) Schematic representation of EGFP under the transcriptional control of an intact *Foxp3* gene. Both, FOXP3 and EGFP will be generated as independent proteins through self-splicing imposed by the T2A sequence. (lower diagram) Sanger sequences of a founder in which in-frame junctions are present between the 5’ and 3’ homology arms of the DNA donor and the *Foxp3* gene as well as the entire donor DNA sequence is conserved with conserved junctions between exon 11-T2A, T2A-EGFP and EGFP-stop sequences. Primers used for the different PCR sequences are outlined.

Rat zygotes were microinjected and were transferred (n=148) into pseudopregnant females. Newborns (n=25) were genotyped by PCR followed by sequencing and one founder was identified as a KI (**Fig. 1B**) whereas other founders (n=4) showed indels **(data not shown).** This founder was mated with a wild-type partner and a *Foxp3-EGFP* rat line was stably generated with a mendelian segregation of the *Foxp3-EGFP* allele (data not shown).

### CD8^+^ and CD4^+^ Treg are the only mononuclear cells that express EGFP

Cytofluorimetric analysis of lymph nodes, thymus, spleen, blood and bone marrow showed that a proportion of TCR^+^CD4^+^ or TCR^+^CD8^+^ cells expressed EGFP that was equivalent to the proportion of FOXP3^+^ cells detected using an anti-FOXP3 antibody (**Fig. 2 and supplementary Fig. 1**). Of note, analysis of homozygote *Foxp3-EGFP* animals showed normal expression of FOXP3 as detected using the anti-FOXP3 antibody and all FOXP3^+^ were also EGFP^+^ **(data not shown)** indicating that the expression of FOXP3 was not impaired by the insertion of the EGFP. NK and B cells as well as monocytes/macrophages did not express EGFP and EGFP was not expressed by cells others than T cells (**data not shown**). A quantitative analysis of CD4^+^EGFP^+^ T cells in thymus and spleen ranged in percentage between 5 and 10% of the cells, respectively (**Table 1**). In absolute numbers, CD4^+^EGFP^+^ T cells ranged between 8×10^4^ and 2.2×10^6^ cells per ml of blood and in the spleen, respectively (**Table 1**). CD8^+^EGFP^+^ T cells represented roughly 10% of the CD4^+^EGFP^+^ cells and ranged between 0.6 and 1.3% of the cells in the thymus and lymph nodes, respectively (**Table 1**). In absolute numbers, CD8^+^EGFP^+^ cells ranged between 0.7 x10^4^ and 0.4×10^6^ cells per ml of blood and in lymph nodes, respectively (**Table 1**).

**Figure 2.**
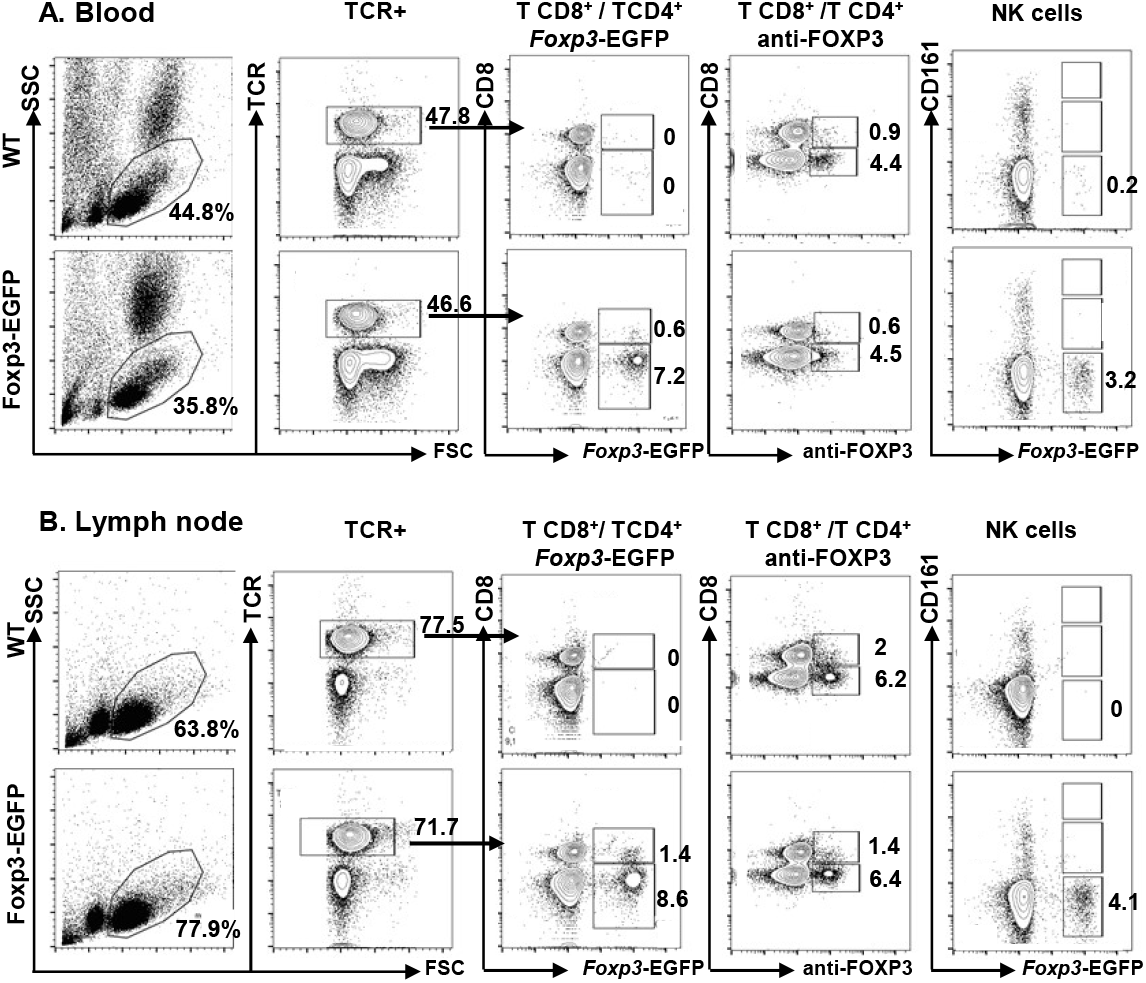
Flow cytometry analysis of peripheral blood lymphocyte and lymph node cells from *Foxp3-EGFP* rats. **A)** Blood and **B)** axillary lymph nodes were harvested from 12-week-old *Foxp3-EGFP* or wild-type (WT) rats and single cell suspensions were gated by SSC and FSC on lymphocytes followed by the identification with mAbs of major cell populations; TCR^+^ cells, TCR^+^CD4^+^, TCR^+^CD8^+^ and CD161^high^ for NK cells. These populations were then analyzed for FOXP3 expression by EGFP expression and by using an anti-FOXP3 mAb. Contour plots from one animal representative of 6 analyzed in the same conditions.

**Table 1.** Quantification of CD4^+^ and CD8^+^ Treg in different immune compartments.

|  | **units** | **thymus** | **LN** | **spleen** | **BM** | **blood/ml** |
| --- | --- | --- | --- | --- | --- | --- |
| **CD4<sup>+</sup>EGFP<sup>+</sup>**<br>(n=5) | % * | 5.2 ± 0.9 | 7.8 ± 1.7 | 10 ± 0.2 | 7.2 ± 3 | 7.9 ± 0.6 |
|  | cellsx10<sup>6</sup> | 2.24 ± 1 | 0.4 ± 0.1 | 2.2 ± 0.4 | 0.19 ± 0.004 | 0.083±0.03 |
| **CD8<sup>+</sup>EGFP<sup>+</sup>**<br>(n=5) | % ** | 0.6 ± 0.15 | 1.3 ± 0.4 | 0.7 ± 0.01 | 0.8 ± 0.45 | 0.7 ± 0.1 |
|  | cellsx10<sup>6</sup> | 0.28 ± 0.2 | 0.4 ± 0.6 | 0.16 ± 0.02 | 0.003 ± 0.002 | 0.007 ± 0.002 |

LN; axillar lymph nodes; BM, bone marrow from 1 femur/animal. \* among TCR<sup>+</sup>CD4<sup>+</sup>;
\*\*among TCR<sup>+</sup>CD8<sup>+</sup>

Thus, *Foxp3-EGFP* rats appear to be a faithful tool to identify and analyze CD4^+^ and CD8^+^ Treg.

### Phenotype and lymphoid organ distribution of TCR^+^CD4+^+^EGFP^+^ and TCR^+^CD8^+^EGFP^+^ Treg

EGFP^+^ cells, both CD4^+^ and CD8^+^ T cells were mostly CD25^+^CD127^low^ cells (**Fig. 3A**), a phenotype largely belonging to CD4^+^FOXP3^+^ Treg in humans, mice and rats (5) but not yet clearly defined for CD8^+^ Treg. EGFP^+^ cells were detected only among CD45RC^low/-^ T cells (**Fig. 3B**), a marker that has been used both for CD4^+^ and CD8^+^ Treg in rats and humans (9, 11, 12). The markers CD25, MHC-II, CD28, and CD44 were more and CD62L less expressed in both CD4^+^ and CD8^+^EGFP^+^ vs. EGFP^-^ cells (**Fig. 3C**). CD5 and CD26 were equally expressed by all cell subtypes **(data not shown).**

**Figure 3.**
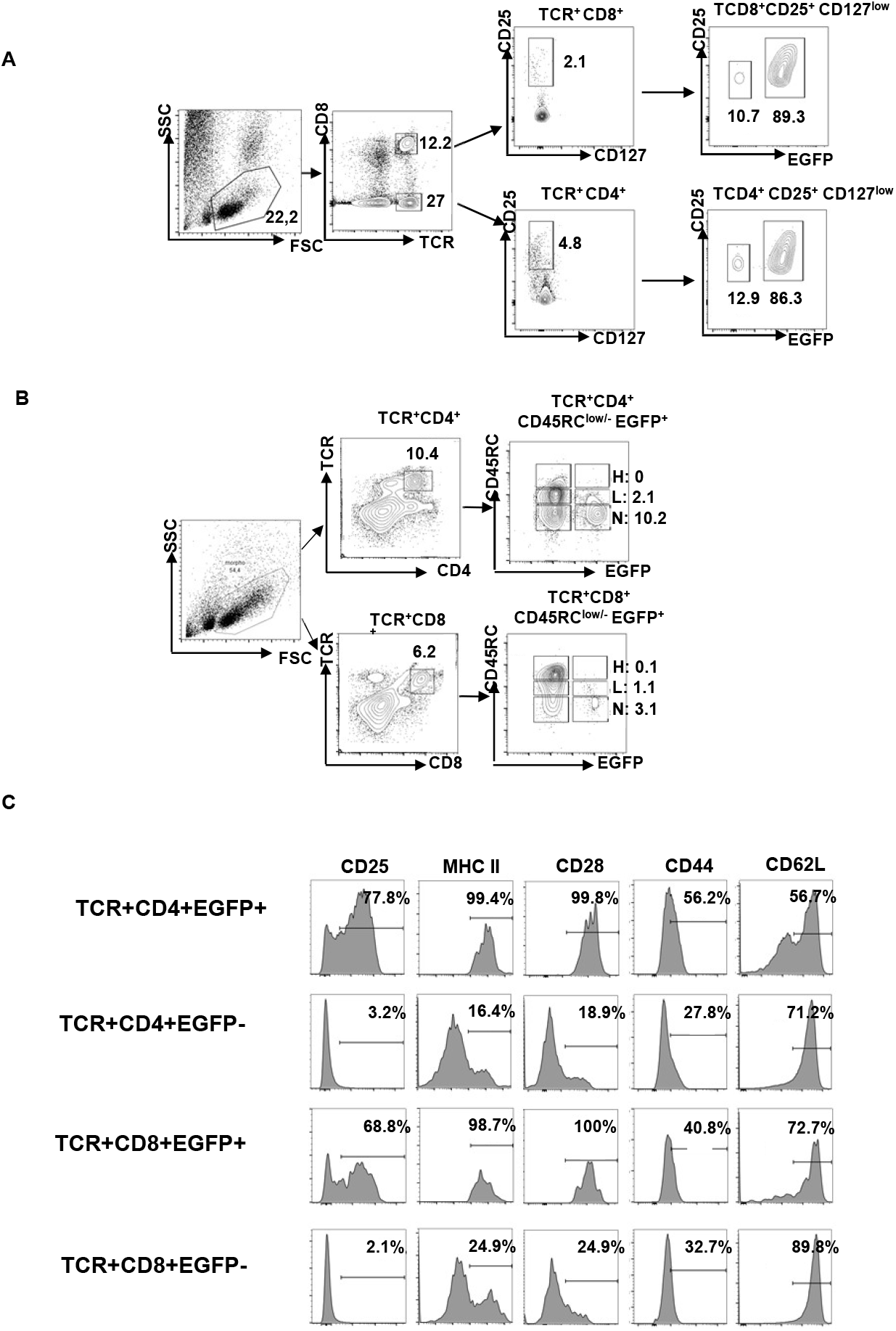
Phenotype of EGFP*^+^* cells. Single cell suspensions were obtained from spleen from a *Foxp3-EGFP* rat, lymphocytes gated by SSC/FSC and analyzed for the phenotype of CD4^+^ and CD8^+^ EGFP^+^ cells. **A)** TCR^+^ cells were labelled with an anti-CD8alpha mAb to identify CD8^+^ and CD8^-^/CD4^+^ and analyzed for CD25 and CD127 expression followed by EGFP detection. **B)** TCR^+^CD4^+^ or TCR^+^CD8^+^ cells were analyzed for CD45RC expression, levels identifying high (H), low (L) and negative (N) cells, and EGFP expression in these populations. **C)** Histograms of TCR^+^CD4^+^EGFP^+^, TCR+CD4+EGFP^-^, TCR+CD8+EGFP+and TCR^+^CD8^+^EGFP^-^ cells analyzed with the indicate mAbs. Percentages above the traits correspond to the number of positive cells above staining with isotype control mAbs. In A, B and C one experiment representative of 5 performed in the same conditions.

The proportion and absolute numbers of TCR^+^CD4^+^EGFP^+^ Treg vs. TCR^+^CD8^+^EGFP^+^ Treg were ∼10-fold higher in thymus, blood, spleen and bone marrow and ∼5-fold higher in lymph nodes (**Table 1**).

We compared the frequency and expression levels of TCR^+^CD4^+^EGFP^+^, TCR^+^CD8^+^EGFP^+^ Treg in rats to the ones in mice also harboring EGFP in the 3’ end of the *Foxp3* gene, and thus as in *Foxp3-EGFP* rats, controlled by the *Foxp3* endogenous promoter (14). The intensity of EGFP expression was comparable **(data not shown).** A quantitative analysis of TCR^+^CD4^+^EGFP^+^ cells in rat vs. mice in different immune compartments showed similar percentage in spleen and bone marrow and higher in thymus, lymph nodes and blood **(Supplementary Table 1)**. Surprisingly, TCR^+^CD8^+^EGFP^+^ cells were increased in rats vs. mice in all compartments, particularly in the thymus, lymph nodes and bone marrow **(Supplementary Table 1)**.

Overall, in rats and mice CD4^+^ Treg are more abundant than CD8^+^ Treg, the phenotype of CD8^+^ Treg resemble the one of CD4^+^Treg when analyzed with a restricted panel of markers and the frequency of CD4^+^Treg and more particularly in CD8^+^Treg was higher in rats vs. mice.

### In vitro suppressive function of EGFP^+^ and EGFP^-^ fractions of CD4+ and CD8+ Treg

We then aimed to confirm the suppressive function of CD4^+^ and CD8^+^ EGFP^+^ Tregs by using a T cell proliferation assay. Present cell surface markers identifying CD8^+^ Tregs are less discriminative than for CD4^+^ Tregs. Since CD8^+^CD45RC^low/-^ and not CD8^+^CD45RC^high^ T cells contain most of the suppressive activity among CD8^+^ cells both in rats (25) and humans\ (7) we sorted CD8^+^CD45RC^low/-^ cells either EGFP^+^ or EGFP^-^ and compared their suppressive function. Although CD4^+^CD25^+^CD127^low^ Tregs contain a very small fraction of EGFP^-^ cells, to keep the same strategy of analysis for CD4^+^ and CD8^+^ Tregs, we sorted CD4^+^CD25^high^CD127^low^ cells EGFP^+^ or EGFP^-^ fractions and compared their suppressive activity. The suppressive activity was confined to the EGFP^+^ fraction for both the CD8^+^CD45RC^low/-^ and CD4^+^CD25^high^CD127^low^ Tregs (**Fig. 4 and supplementary Figure 2**). Thus, isolation of viable CD4^+^ and CD8^+^ Treg by their expression of EGFP allows to explore their suppressive function.

**Figure 4.**
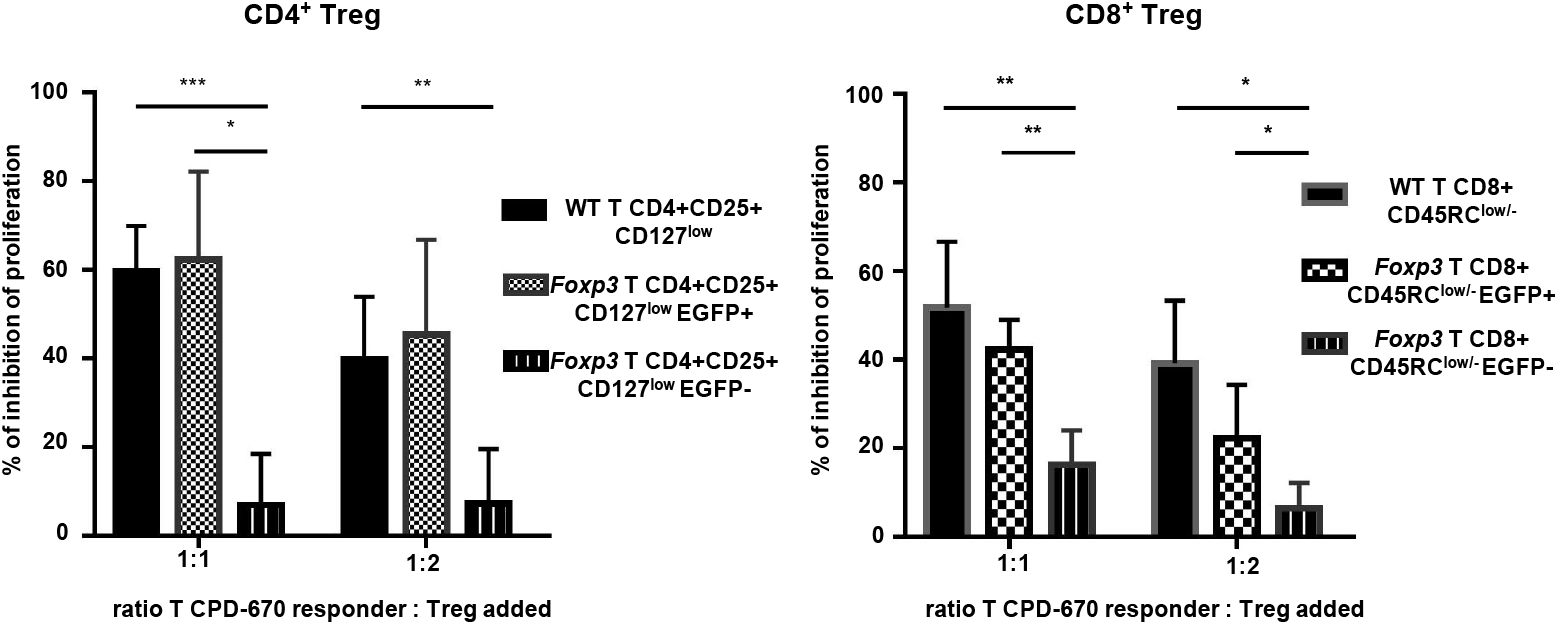
Suppressive activity of TCR^+^CD4^+^EGFP^+^ and TCR^+^CD8^+^EGFP^+^ cells in an MLR. An MLR was performed by co-culturing in a 1:1 ratio spleen CD4^+^CD25^-^ Tconv cells from SD/Crl rats (MHC haplotype u) with enriched spleen APCs from Lewis 1A (MHC haplotype a) rats. Spleen Tregs from wild-type (WT) or *Foxp3-EGFP* SD/Crl rats were sorted based on their surface phenotype using previously defined markers. For CD4^+^ Tregs, TCR^+^CD4^+^CD25^+^CD127^low/-^ and for CD8^+^ Tregs, TCR^+^CD8^+^CD45RC^low/-^. CD4^+^ and CD8^+^ Tregs were then sorted based on the expression or not of EGFP. The four populations of Tregs, CD4^+^EGFP^+^ and EGFP^-^ as well as CD8^+^EGFP^+^ and EGFP^-^ were added to the MLR in a 1:1 or 1:2 Tregs to CDP-670 labelled Tconv cells. Values are expressed as mean % +/− SEM of suppression normalized to MLR proliferation in the absence of Tregs (0 % inhibition of proliferation). n=4. *p<0.05, **p<0.01, **p<0.001.

### Transcriptomic analyses of natural and induced EGFP^+^ CD8^+^ and CD4^+^ Treg

To gain further insight into the molecular properties of CD4^+^ and CD8^+^ Treg we sorted CD25^high^CD127^low^CD4^+^EGFP^+^, CD25^high^CD127^low^CD4^+^EGFP^-^, CD45RC^low/-^CD8^+^EGFP^+^ and CD45RC^low/-^CD8^+^EGFP^-^ cells and compared their transcriptomic profiles. Principal components analysis (PCA) showed that CD4^+^EGFP^+^ vs. CD4^+^EGFP^-^ (**Fig. 5A left)** and CD8^+^EGFP^+^ vs. CD8^+^EGFP^-^ T cells (**Fig. 5A middle)** clustered separately. Notably, there was no clear distinction between the CD4^+^EGFP^+^ and CD8^+^EGFP^+^ T cells (**Fig. 5A right)**. Comparison gene expression revealed multiple differentially expressed genes (DEG) between the different cell types (FDR Q-value <0.05) (**Fig. 5B**). When comparing CD4^+^EGFP^+^ vs. CD4^+^EGFP^-^ cells, CD4^+^EGFP^+^ cells had 309 genes with higher expression. Among them were several Treg specific genes (eg: *Foxp3, Ikzf2, Tox, Lrrc32(GARP))*. Whereas CD4^+^EGFP^-^ cells had 290 genes upregulated with some genes with a CD4^+^ Th cell specific (eg: *Nkg7, Gzmb, Cd40lg*…) (**Fig. 5B left, supplementary table 2)**. Comparison of CD8^+^EGFP^+^ vs. CD8^+^EGFP^-^ cell, revealed that CD8^+^EGFP^+^ cells had 373 genes with higher expression, again with genes that are specific to Treg such as (eg: *Foxp3, Lrrc32*(GARP)*, Il2ra*) but others acts as regulators of CD8^+^ (eg: *C1qa* and *C1qc*) whereas CD8^+^EGFP^-^ cells 363 genes upregulated, such as genes critical for the cytotoxic function of effector *CD8*^+^ T (eg : *Gzmk, Gzmm..)* (**Fig. 5B middle, supplementary table 3)**. When comparing CD4^+^EGFP^+^ vs. CD8^+^EGFP^+^ cells, CD4^+^EGFP^+^ cells upregulated 322 genes, such as *Gm2a, Smpdl3a* and *Nrp1.* CD8^+^EGFP^+^ upregulated 242 genes, such as *Lag3, Np4, C1qa,* and *Tnfrsf18* (**Fig. 5B right, supplementary table 4)**.

**Figure 5.**
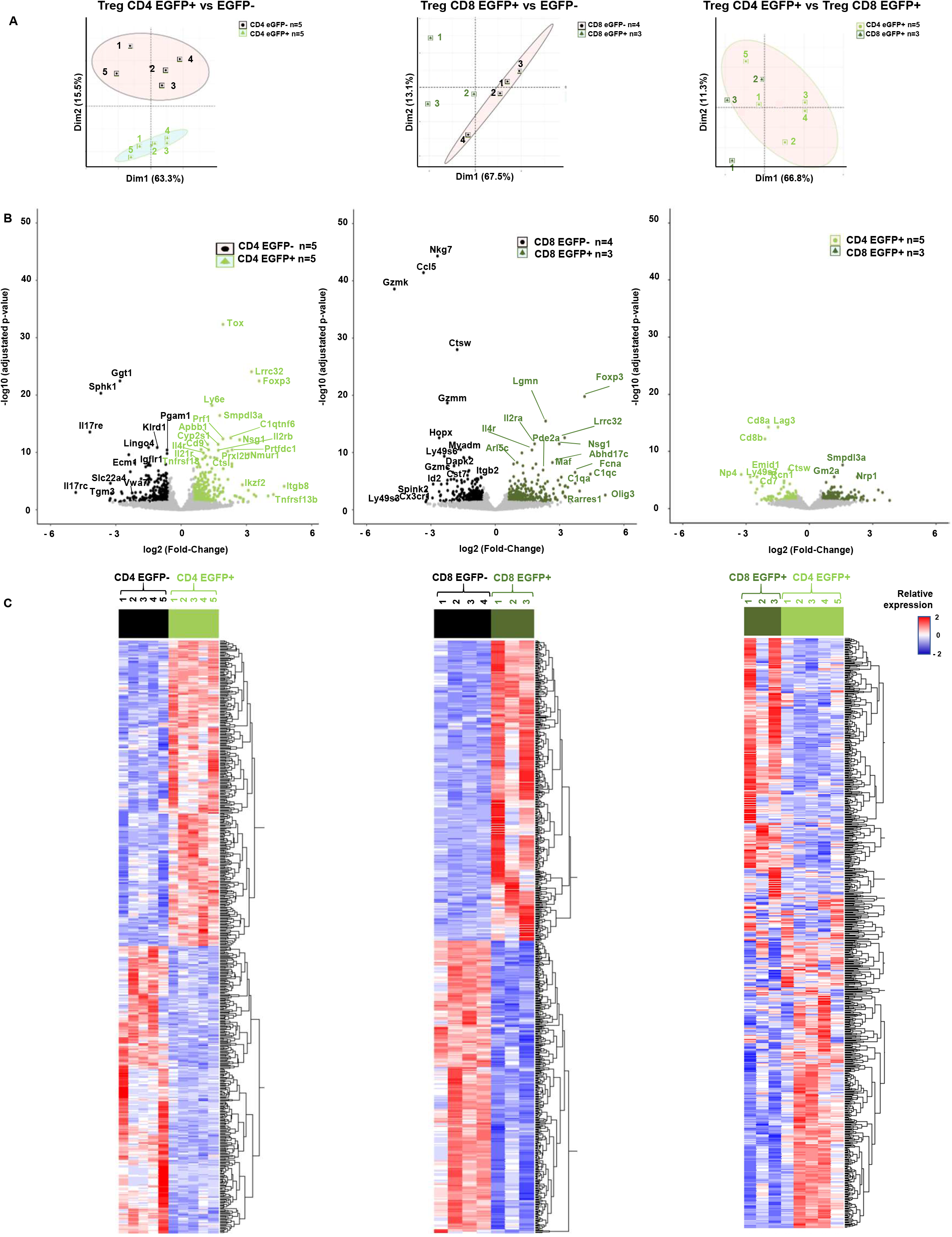
RNAseq analyses of EGFP^+^ and EGFP^-^ within CD4^+^ and CD8^+^ T cells. TCR^+^CD25^high^CD127^low^CD4^+^ and TCR^+^CD45RC^low/-^ EGFP^+^ and EGFP^-^ T cells were cell sorted and RNAseq analyses were performed comparing CD4^+^EGFP^+^ vs. CD4^+^EGFP^-^ cells (left column), and CD8^+^EGFP^+^ vs. CD8^+^EGFP^-^ cells (middle column) and CD8^+^EGFP^+^ vs. CD4^+^EGFP^+^ cells right column. **A.** principal components analysis (PCA) of all samples and of expressed genes. **B.** Volcano plots. (Left) Light green indicates overexpression in CD4^+^EGFP^+^ vs. CD4^+^EGFP^-^ cells and black of CD4^-^EGFP^+^ vs. CD4^+^EGFP^+^ cells. (middle) Dark green indicates overexpression in CD8^+^EGFP^+^ vs. CD8^+^EGFP^-^ cells and black of CD8^-^ EGFP^+^ vs. CD8^+^EGFP^+^ cells. (right) Light green indicates overexpression in CD4^+^EGFP^+^ vs. CD8^+^EGFP^+^ cells and dark green of CD8^-^EGFP^+^ vs. CD4^+^EGFP^+^ cells. **C.** Heatmaps of all differentially expressed genes. Low expression levels are in blue, mean expression levels are in white and high expression levels are in red. Each sample number is depicted on the top.

Heatmap analysis of DEG grouped together with different transcriptomic profiles all CD4^+^EGFP^+^ vs. CD4^+^EGFP^-^ cells, CD8^+^EGFP^+^ vs. CD8^+^EGFP^-^ cells, as well as CD4^+^EGFP^+^ cells vs. CD8^+^EGFP^+^ cells (**Fig. 5 C**). Among these DEG many were genes of the immune system and for CD4^+^EGFP^+^ cells with a fold change of >1-fold there were already identified key genes such as some described above as well as *Itgb8, Cd80* and *Tnfrsf9* **(Supplementary Table 2**) (26–30).

For CD8^+^EGFP^+^ cells with a fold change of >1-fold there were genes shared with CD4^+^EGFP^+^ Treg such as some described above as well as *Il4r, Fcna* and *Art2b* as well as others previously described in human or mouse CD4^+^ Treg, such as *Il2ra, Lgmn* (31) and *Tnfrsf9* **(Supplementary Table 3).**

We then compared upregulated genes with immune functions in CD4^+^EGFP^+^ vs. CD8^+^EGFP^+^ Treg **(Supplementary Table 4).** In CD4^+^EGFP^+^ Treg we observed increase expression of genes such as *MHC-II (LOC688090), Lrrc32, Ccr6, Selplg and Tnfrsf4* **(Supplementary Table 4)**. In CD8^+^EGFP^+^ Treg we observed upregulation of genes with immune function not previously linked to Treg, such as *Fcna*, *Hmox1*, *Spic* and complement pathway genes (*C1qc, C1qa, and C1qb)* **(Supplementary Table 4)**. Further analysis of expression levels of selected immune genes in individual samples of the four populations of cells showed that some were upregulated in both CD4^+^EGFP^+^ and CD8^+^EGFP^+^ populations vs. the EGFP^-^ counterparts, such as *Foxp3, Lrrc32, Tnfrsf1b, Tnfrsf9, Il2ra, Tox* and *Il4r* whereas others were only upregulated in CD4^+^EGFP^+^ Treg, such as *Stap2* and *Igtb8* or in CD8^+^EGFP^+^ Treg, such as *Timp1* and *Erc1* **(Supplementary Fig. 3A).**

Based on the genes differently expressed and the biological pathways in which they are involved we determined 4 principal processes with different pathways and with differential involvement for each cell type: immune cells activation, proliferation and adhesion as well as a miscellaneous one **(Supplementary fig 3 B)**. CD4^+^EGFP^+^ vs. CD4^+^EGFP^-^ cells, expressed genes involved in immune cell activation and adhesion but not in immune cell proliferation and several of these pathways were shared with CD8^+^EGFP^+^ Treg. CD8^+^EGFP^+^ vs. CD8^+^EGFP^-^ cells also expressed genes involved in immune cell proliferation or its regulation. CD4^+^EGFP^+^ vs. CD8^+^EGFP^+^ showed genes involved in the miscellaneous pathways, such as Th1 and Th2 cell differentiation and regulation of inflammatory responses.

Venn analyses of all genes upregulated by CD4^+^EGFP^+^ vs. CD8^+^EGFP^+^ Treg vs. their EGFP^-^ counterparts as well as between the two Treg populations allowed to define genes that were unique and common among them **(Supplementary Fig. 3C)**. The genes up and down regulated in the Venn diagrams are listed in **Supplementary Table 5 and 6,** respectively.

We finally confirmed at the protein level some of the genes that were upregulated at the RNAseq level such as for CD25/*Il2ra* (**Figure 3 and Supplementary Fig. 3A, respectively)**, as well as CD44/*Cd44* and ICOS/*Icos* **(Supplementary Fig. 4).**

Altogether, the transcriptome of CD4^+^EGFP^+^ and CD8^+^EGFP^+^ Treg showed common genes previously described for CD4^+^ Treg but also many others that differ and that may allow to define a CD8+ Treg transcriptomic signature.

### Analysis of in vitro generated EGFP^+^ induced Tregs

We tested whether EGFP^-^ Tconv cells from *Foxp3-EGFP* rats could be converted to EGFP^+^ induced Tregs. To this end, CD4^+^CD25^-^ and CD8^+^CD45RC^low/-^EGFP^-^ T cells were isolated and cultured in the presence of anti-CD3 and CD28 mAbs, IL-2 and TGFbeta. Stimulation with IL-2 and in the absence of TGFbeta has been shown to induce FOXP3 in T cells that were FOXP3^-^ in humans (32), likely as a marker of T cell activation, but not in mice (33) whereas in the presence of IL-2 and TGFbeta FOXP3 is induced in both species (32, 33). The induction of FOXP3 in rat T cells has not been clearly addressed. In the presence of anti-CD3 and anti-CD28 and TGFbeta alone, CD4^+^CD25^-^EGFP^-^ and CD8^+^CD45RC^low^EGFP^-^ T cells converted to 19.7 % and 7.9 % of EGFP^+^ induced Tregs, respectively, and the MFI expression of EGFP was very similar than the one of EGFP^+^ natural Tregs (for CD4^+^EGFP^+^: 9711 vs. 8394, respectively and for CD8^+^EGFP^+^ 11000 vs. 13000, respectively) (**Fig. 6 A**). The culture with IL-2 alone or IL-2 and TGF beta did not further increase the percentage of EGFP+ cells compared to culture medium alone or TGF-beta alone, respectively (**Fig. 6 A**).

**Figure 6.**
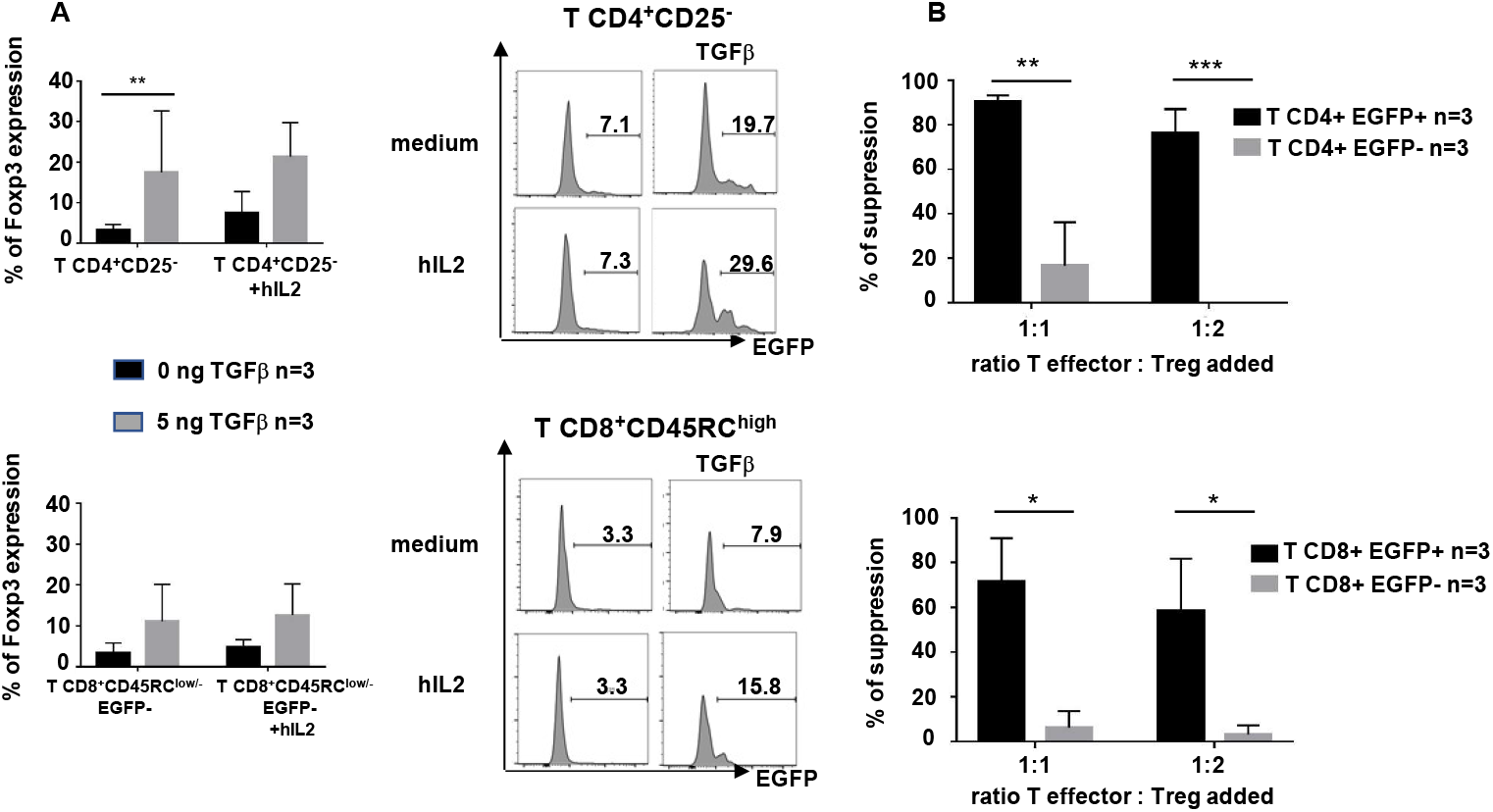
Conversion of EGFP^-^ T cells in EGFP^+^ induced Treg. Spleen cells from *Foxp3*-EGFP animals were used to sort Tconv CD4^+^CD25^-^ Treg and T CD8^+^CD45RC^low/-^EGFP^-^ cells, labelled them with CPD-670 and cultured for 3 days in the presence of anti-CD3/CD28 mAbs, IL-2 and in the absence or presence of hTGFbeta. **A.** Cells that proliferated (CPD- 670low/neg) were then analyzed for EGFP expression. Left graphs show mean +/− SEM of 3 different experiments and right histograms show one representative analysis. **B.** Suppressive assays using EGFP^+^ (CD4^+^ or CD8^+)^ vs. EGFP^-^ cells (CD4^+^ or CD8^+^) from the same cultures at different ratios with CD4^+^ Tconv responder cells. n=3, mean +/− SEM.

We then evaluated the suppressive activity of the induced EGFP^+^ Tregs from the condition in the presence of TGFbeta and IL-2 vs. the T cells from the same cultures that were EGFP^-^. Induced CD4^+^ and CD8^+^ EGFP^+^ Tregs were highly suppressive of MLR proliferation and significantly more than EGFP^-^ cells from the same cultures that only suppressed MLRs marginally (**Fig. 6 B**).

In conclusion, the use of *Foxp3-EGFP* rats allowed to define in vitro conditions of generation of induced Treg and to demonstrate that both T CD4^+^ and CD8^+^ lineages could generate induced Treg.

### IL-2 induced expansion of EGFP^+^ Tregs in vivo

In vivo injection of low doses of IL-2 has been shown to preferentially expand CD4^+^ and CD8^+^ FOXP3^+^ Treg in mice (34), nonhuman primates (35) and humans (36). In rats, CD4^+^FOXP3^+^ Treg were increased upon administration of IL-2 and CD8^+^FOXP3^+^ Treg were not described (37). We aim to define whether we could detect expansion of CD4^+^ and CD8^+^ EGFP^+^ Treg upon IL-2 in vivo injection. Injection of IL-2 resulted in a significant expansion of EGFP^+^ cells as compared to untreated rats in both the CD4^+^ (from 5.6 ± 1.4 to 17.1 ± 0.6) and CD8^+^ (from 2.2 ± 0.4 to 9.8 ± 0.9) T compartments as well as a decrease of the reciprocal EGFP^-^ compartments whereas NK cells remained EGFP^-^ (**Fig. 7A**). In blood of IL-2-treated animals compared to controls increased absolute numbers of T CD4^+^EGFP^+^ (19-fold) and CD8^+^EGFP^+^ (31-fold) cells showed a higher increase vs. T CD4^+^EGFP^-^ (4.4-fold) and CD8^+^EGFP^-^ (14-fold) cells and NK cells (10-fold) **(supplementary Table 7).** The proportions of T CD4^+^ and CD8^+^ EGFP^+^ cells were also increased in the spleen **(supplementary Fig. 5).** Phosphorylated Stat5 (**Fig. 7B**) and CD25 (**Fig. 7C**) were increased in EGFP^+^ cells but also in EGFP^-^ T cells.

**Figure 7.**
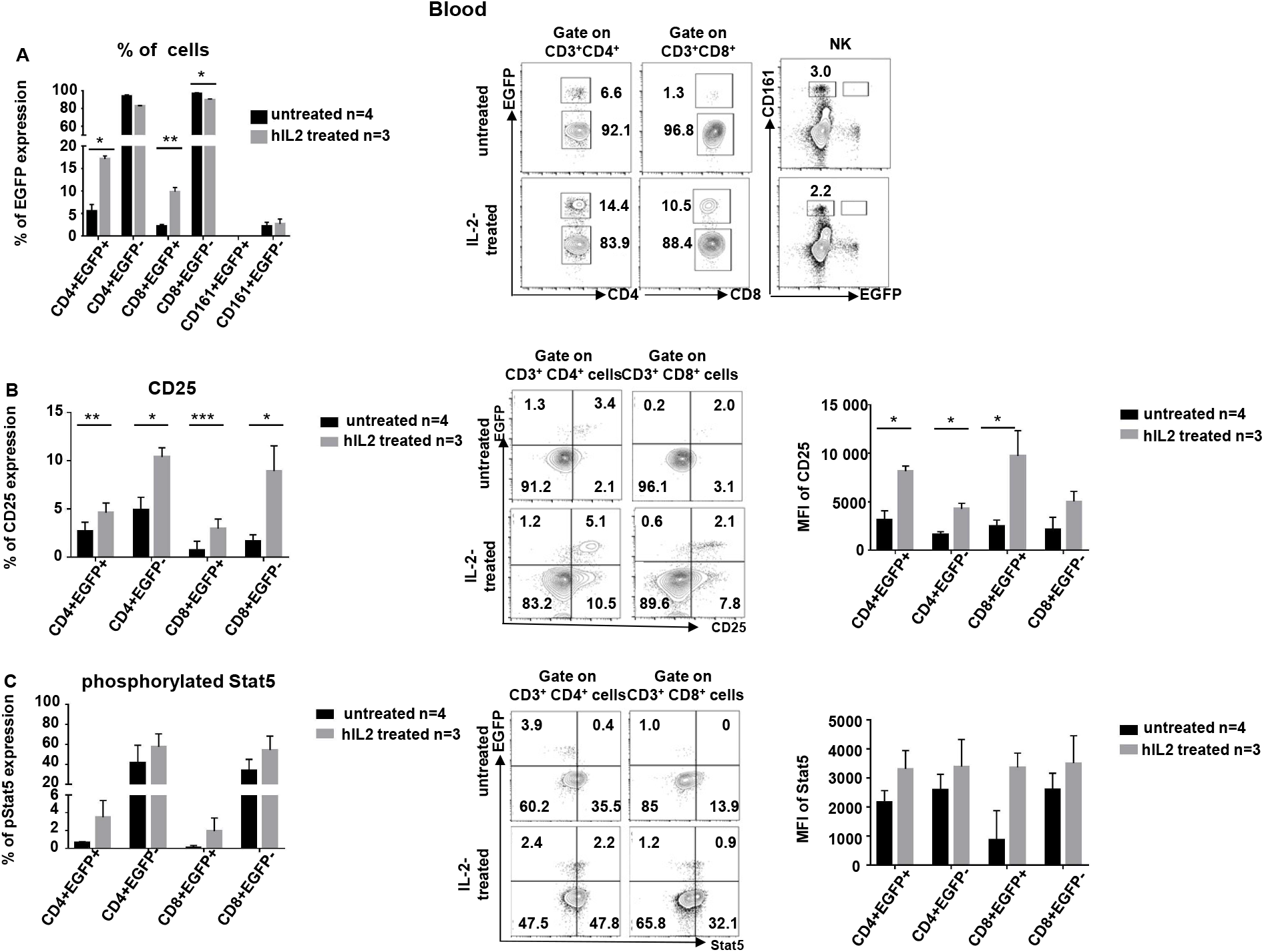
IL-2 induce expansion of EGFP^+^ Tregs in vivo. *Foxp3*-EGFP rats received or not hIL-2 and the percent of the indicated cell populations was analyzed in blood. **A.** Percentages of cell EGFP^+^ cells in the indicated cell populations of IL-2-treated and untreated rats. Left graph shows mean +/− SEM, n=3, and right contour plots show a representative animal. **B.** Percentage and MFI (left and middle histograms, respectively) of pStat5^+^ cells in the indicated cell populations in IL-2-treated and untreated rats. (mean +/− SEM, n=3). Middle contour plots show a representative animal. **C.** Percentage and MFI (left and middle histograms, respectively) of CD25^+^ cells in the indicated cell populations of IL-2-treated and untreated rats. (mean +/− SEM, n=3). Middle contour plots show a representative animal.

Thus, *Foxp3-EGFP* rats allow to detect increases in CD4^+^ and CD8^+^ Treg induced by IL-2.

## Discussion

*Foxp3-EGFP* rats were generated using a strategy in which *Foxp3* gene drives the expression of EGFP placed in the 3’ end of the gene. This strategy preserves Foxp3 expression of the allele in which the transgene is inserted avoiding decrease expression in heterozygous animals and also allows the use of homozygous animals to more efficiently maintain the mutated rat line. EGFP was exclusively expressed by CD4^+^ and CD8^+^ T cells and in proportions like the ones defined using anti-FOXP3 antibodies and thus EGFP was strictly restricted to subsets of the T cell lineage. CD4^+^EGFP^+^ Treg were CD25^high^CD127^low/-^ CD45RC^low/-^, consistent with the what has already been previously described in rats, humans and mice. All CD8^+^EGFP^+^ Treg were also CD25^high^CD127^low/-^CD45RC^low/-^. Quantitative analysis of *Foxp3-EGFP* rats showed that CD4^+^EGFP^+^ Treg were 5-10-fold more abundant than CD8^+^EGFP^+^ Treg.

Genetic labelling of *Foxp3*-expressing cells preserving *Foxp3* expression has been previously used in some mouse models (14, 16, 18–20). Other mouse *Foxp3-eGFP* models generated a knockout of the allele in which GFP was inserted (15, 17). In some of these mouse models, CD8^+^GFP^+^ Treg were described with frequencies ranging between 5 to-100 fold lower than CD4^+^GFP^+^ Treg (15–18), whereas in the others they were not described. Analysis in our experimental conditions of CD8^+^GFP^+^ Treg in one of this mouse models (14) showed 9 to 130-fold more CD4^+^EGFP^+^ Treg vs. CD8^+^EGFP^+^ Treg. To the best of our knowledge there are no other mammalian species with transgenic lines allowing to define FOXP3^+^ cell lineages.

CD4^+^ Treg defined by the usual CD25^high^ and CD127^low/-^ markers showed a majority of EGFP^+^ cells but also the presence of EGFP^-^ cells. Analysis of their suppressive activity revealed that >90% was confined to the EGFP^+^ fraction of cells. Similarly, CD8^+^ Treg defined using CD45RC^low/-^ as a marker also showed EGFP^+^ and EGFP^-^ cells and most of the suppressive activity was observed in the EGFP^+^ vs. EGFP^-^ fraction.

Transcriptomic analysis revealed for CD4^+^EGFP^+^ Treg the upregulation of genes previously described in mouse and/or human CD4^+^ Treg as important for their differentiation, suppressive function and oligodendrocyte regeneration, such as *Itgb8, Tnfrsf13b, Foxp3, Ikzf2, Ccn3, Il1rl1, Lrrc32, Cd80, Tnfrsf9, Gata3, Il2ra, Il2rb, Tnfrsf1b* (TNFR2 or CD120b), *Tnfrsf4* (OX40 or CD134) *and Tnfrsf18* (GITR or CD357) (26-30, 38-40). CD4^+^EGFP^+^ Treg also highly upregulated genes coding with immune function not previously described for CD4^+^ Treg, such as *Stap2* (41), *Tox* (42) and *C1qtnf6* a soluble inhibitor of the alternative complement activation pathway, for which further work is needed to define their functional role. Transcriptomic analysis of CD8^+^EGFP^+^ Treg is to the best of our knowledge the first one to be described. CD8^+^EGFP^+^ Treg shared some genes previously described in CD4^+^EGFP^+^ Treg, such as *Lrrc32, Il2ra, Tnfrsf1b, C1qtnf6* and *Tnfrsf9,* but not other such as *Ctla4* and *Gata3,* (26-31, 38-40). CD8^+^EGFP^+^ Treg expressed immune genes that need to be analyzed for their functional role in Treg, such as complement genes (*C1qc, C1qa, C1qb, Cfd*) and at the same time *C1qtnf6* (a secreted molecule that inhibits complement activation), *Fcna* (a secreted molecule with elastin-binding activity), *Hmox1* (enzyme that degrades heme), *Spic* (an enhancer of transcription in lymphoid cells), and *Vipr2* (one of the receptors for vaso intestinal peptide).

Since there is need to identify cell membrane markers that could allow to purify CD45RC^low/-^ CD8^+^FOXP3^+^ Treg, several genes that code for membrane markers that were upregulated in CD45RC^low/-^CD8^+^EGFP^+^ Treg or in CD45RC^low/-^CD8^+^EGFP^-^ Treg were identified that could be used to identify and isolate CD8^+^EGFP^+^ Treg by positive or negative selection, respectively. Genes upregulated in CD8^+^EGFP^+^ Treg included *Vcam1 (CD106), Cdh1 (CD324), CD163, Mrc1 (CD206), Trpm2, Kdr (CD309), FcmR, Itgb5, Il9r (CD129), Il2ra (CD25), Cd79b, Ly49s7, Jag1 (CD339), Il6st (CD130), IL4r (CD124), Vipr2, Tfrc (CD71), Il6r (CD126)* and *Tnfrsf1b (TNFR2).* In CD8^+^EGFP^-^ Treg genes encoding for cell membrane molecules that were upregulated included *Ly49s3, Nkg7, Ly49s3, Fcnb, several RT1 (MHC-II) molecules, Nrc1 (CD335), Klrk1 (CD314), CD38, CD59 and CD174*. For this, the RNAseq data need to be confirmed by protein cytometry analyses, as done for CD25, CD44 and ICOS, since not only mRNA levels may not translate into higher protein levels but also because the RNAseq analysis on whole cell populations do not allow to conclude whether it is a large or a small fraction of the cells that upregulate a given gene. Thus, these cell membrane markers could potentially be used to identify and purify CD8^+^ nTreg in wild-type rats and even in other species. On this regard, human CD8^+^CD45RC^low/-^ cells contain the CD8^+^ Treg suppressive activity but further enrichment of cells positives for CD25 or negatives for CD127 did not increase the suppressive activity compared to the total CD8^+^CD45RC^low/-^ population (7). This suggest a difference between human and rat nCD8^+^ Treg for these markers but human CD8^+^CD45RC^low/-^FOXP3^+^ cells did express higher levels of GITR than CD8^+^CD45RC^low/-^FOXP3^-^ cells and were more suppressive than the GITR^-^ fraction of CD8^+^CD45RC^low/-^ cells (7) showing a concordance with the transcriptomic data presented in this manuscript.

Despite that most suppressive activity was confined in nTreg to the rat EGFP^+^ fraction for both CD4^+^ and CD8^+^ T cells, in vitro and in vivo iTreg have been described (1, 3, 4, 6, 7). In vitro culture Tconv CD4^+^CD25^-^ EGFP^-^ or CD8^+^CD45RC^low/-^GFP^-^ cells in the presence of TGFbeta and IL-2 but not of IL-2 alone resulted in the induction of EGFP in a fraction of cells. The suppressive activity of the cultured cells was restricted to the EGFP^+^ fraction of cells in both CD4^+^ and CD8^+^ T cell compartments and these are thus truly iTreg. This suggests that *Foxp3-EGFP* rats could be used to analyze iTreg in different situations in vitro and in vivo. Generation of FOXP3^+^ cells from CD4^+^FOXP3^-^ in humans can be obtained with IL-2 in the absence of TGFbeta but these cells may not be truly Treg since expression of FOXP3 by human cells can be a marker of activation (33). On the contrary, IL-2 alone in mice does not induces FOXP3^+^ cells (32). In both species, mice and human, TGFbeta induces the expression of FOXP3 (32). Thus, induction of expression of FOXP3 in rat Teg cells could be similar to the one observed in mice and different from the one in human Treg but this needs further experiments with additional stimuli.

Low dose IL-2 treatment was shown to preferentially expand both CD4+ and CD8+ FOXP3+ Treg in mice (34, 43), non-human primates (35) and humans (36). In mice, the doses used ranged between 5×10^3^ and 5×10^4^ IU/g of body weight (34, 43). The dose of IL-2 used in this study also (10^4^ IU/g of body weight) was the same used in a previous study in rats that showed increase in CD4+ Treg but in which CD8+ Treg were not described (37). Although with this dose of IL-2 we observed a preferential expansion and activation of both CD4^+^ and CD8^+^ EGFP^+^ Treg, we also observed increased CD4^+^ and CD8^+^ Tconv and NK cells and thus future experiments could analyze lower doses to define a more restricted effect on Treg. It should be noted that doses of 2×10^3^ IU/rat in HLA-B27 with spondyloarthropathy had a weak effect on CD4+ Treg induction (44).

Similar percentages of CD4^+^FOXP3^+^ Treg have been described in different rat strains and CD4^+^ and CD8^+^ Treg in rats have been studied in a variety of pathophysiological situations (5, 6). Although the role of CD8^+^ Treg is less studied than for CD4^+^ Treg, there is solid evidence that this subset of Treg plays an important role in immune responses not only in rats but also in all other species analyzed (4, 7, 9, 10, 45).

## Conclusions

*Foxp3-EGFP* rats are a useful model to identify CD4^+^ and CD8^+^ Treg and to define new molecules expressed in by CD4^+^ but particularly CD8^+^ Treg that will allow to better define their phenotype and function.

The immune system of rats have several characteristics that make them more similar to humans than mice (21) and the recent generation of many gene edited rats (22) will allow to cross *Foxp3-EGFP* rats with other mutated strains to analyze the role of Treg in these models.

## Materials and Methods

### Animals

Wild type Sprague-Dawley (SD/Crl) rats were from Charles River (L’Arbresle, France). *Foxp3-GFP* founder reporter mice were kindly provided by Bernard Malissen (14). All the animal care and procedures performed in this study were approved by the Animal Experimentation Ethics Committee of the Pays de la Loire region, France, in accordance with the guidelines from the French National Research Council for the Care and Use of Laboratory Animals (Permit Number : Apafis 692). All efforts were made to minimize suffering. The rats were housed in a controlled environment (temperature 21±1°C, 12-h light/dark cycle).

### Generation and genotyping of *Foxp3-EGFP* animals

Several CRISPR sgRNA sequences cleaving immediately 3’ of the stop codon of *Foxp3* were designed using the CRISPOR software. The in vitro transcribed sgRNA were purified and one selected for highest cleavage in the rat C6 cell line, as previously described in detail (46, 47). Fertilized one-cell stage embryos were collected for subsequent microinjection using a previously published procedure (48). Briefly, a mixture of Cas9 protein (50 ng/μl), sgRNA (GCAGGGGTTGGAGCACTTGC) (10 ng/μl) and donor DNA (2 ng/μl) encoding the 2A self-cleaving peptide and EGFP sequences flanked by homology arms (1 kb each) 5’ and 3’ from the DNA cleavage point (**Fig. 1A**) was microinjected both into the male pronucleus and into the cytoplasm of fertilized one-cell stage embryos. Microinjected zygotes were maintained under 5% CO2 at 37°C for 2h. Surviving embryos were implanted on the same day in the oviduct of pseudo-pregnant females (0.5 dpc) and allowed to develop to full term. For genotyping rats, DNA from tail biopsy from 8- to 10-day-old rats were digested in 500 _μ_L of tissue digestion buffer (Tris–HCl 0.1 mol/L pH 8.3, EDTA 5 mmol/L, SDS 0.2%, NaCl 0.2 mol/L, PK 100 _μ_g/mL) in a 1.5 mL tube at 56°C overnight and genotyped by PCR using the following primers: *Foxp3*-up; 5’- AAC CTG GGG CTA AAT GTG TG -3’; *Foxp3*-low; 5’- TAG GGT TTG GGT TGA GTC CA -3’; *EGFP*-up; 5’- CCT CGT GAC CAC CCT GAC CT-3’; *EGFP* -low; 5’- TCC ATG CCG AGA GTG ATC CC -3’. PCR amplicons were analyzed by capillary electrophoresis as described (49), followed by Sanger sequencing in *Foxp3*-mutated founders.

### Cytofluorimetry and antibodies

Single-cell suspensions from the spleen, thymus, bone marrow and lymph nodes were prepared as described previously (50). Cell suspensions were analyzed using antibodies (antibodies used are listed in supplementary Table 8). The incubation period was 30 min at 4°C, and the analysis was performed with a FACSVerse system (BD Biosciences, Franklin Lakes, NJ) and FlowJo software (Tree Star, Ashland, OR).

### In vitro suppressive assays

Suppressive assays were performed as previously described in detail (25, 45). Briefly, CD4^+^CD25^−^ naive T cells, CD4^+^CD25^+^CD127^low^ and CD8^+^CD45RC^low/-^ T cells expressing or not FOXP3 were FACS-sorted from *Foxp3-EGFP* or WT rats (haplotype RT-1u). CD4^+^CD25^−^ naive T cells were CPD-670-labeled. Naïve T cells and CD4^+^CD25^+^CD127^low^ (both labeled with CPD-450) and CD8^+^CD45RC^low/-^ T cells were co-cultured at different ratios with spleen DCs from Lew.1A rats (haplotype RT-1a) for 6 d at 37C° in 5% CO2. Proliferation of CPD 670-labeled CD4^+^CD25^−^ naive T cells was analyzed by gating on TCR^+^CD4^+^ cells among live cells and analyzing CPD-670 signal dilution.

### Transcriptomic analysis of CD8^+^ and CD4^+^ EGFP^+^ cells

We isolated TCR^+^CD25^high^CD127^low^CD4^+^EGFP^+^, TCR^+^CD25^high^CD127^low^CD4^+^EGFP^-^, TCR^+^CD45RC^low/-^CD8^+^EGFP^+^ and TCR^+^CD45RC^low/-^CD8^+^EGFP^-^ cells and performed total RNAseq by performing 3’Digital Gene Expression (3’DGE) RNA-sequencing. A RNeasy-Mini Kit (Qiagen) were used to isolate total RNA that was then processed for RNA sequencing. Protocol of 3’digital gene expression (3’DGE) RNA-sequencing was performed as previously described (7). Briefly, the libraries were prepared from 10ng of total RNA. The mRNA poly(A) tail is tagged with well specific barcodes and unique molecular identifier (UMI) and flanked by universal sequencing adapters during the template-switching reverse transcriptions.

Barcoded cDNAs from multiple samples are then pooled, amplified and tagmented using a transposon-fragmentation approach which enriches for 3’ends of cDNA. A library of 350-800bp is sequenced on a S1 flowcell on a NovaSeq 6000 (Illumina). Digital gene expression (DGE) profiles are generated by counting for each sample, the number of unique UMIs associated with each RefSeq genes. Counts matrix was normalized by a linear model with the R package DESeq2 (51). Differential gene expression was determined Wald test and corrected with the False Discovery Rate (FDR) multiple testing method. Genes were considered differentially expressed if the FDR <= 0.05 and the absolute value of log_2_(Fold Change) greater than 0.5. Heatmaps were generated with the R package gplots (https://cran.r-project.org/web/packages/gplots/index.html) Volcano plots were drawn by plotting -log10 (FDR) in function of log2(Fold-Change).

Genes were highlighted when absolute value of log2(Fold-Change) was superior to 1 and FDR <= to 0.05. Principal component analyses (PCA) were performed on all expressed genes with the R package FactoMineR (52). Both pathways from the Kyoto Encyclopedia of genes and genomes (KEGG)(53) as well as gene ontologies (GO) (54) were tested for enrichment among differentially expressed genes with the R package cluster Profiler (55). The significant GO terms and their enrichment scores were filtered with a corrected p-value <= 0.05 (Benjamini-Hochberg method).

### In vitro generated EGFP^+^ induced Tregs

EGFP^-^CD4^+^CD25^-^ and EGFP^-^CD8^+^CD45RC^low/-^ T cells were FACS-sorted from *Foxp3*-*EGFP* rats, labeled with CPD 670 and cultured for 3 days in the presence of anti-CD3 and anti-CD28 MAbs with different doses of hTGFbeta (0, 5, 20 μg/ml) and with or without 100 IU of hIL-2 at 37°C in 5% CO2. Proliferation of CPD-labeled T cells measured by dilution of the CPD signal and EGFP expression were analyzed using a FACSVerse system (BD Biosciences, Franklin Lakes, NJ) and FlowJo software (Tree Star, Ashland, OR). EGFP^+^ CD4^+^ and CD8^+^ cells as well as EGFP^-^ CD4^+^ and CD8^+^ cells were sorted at the end of the culture period and used in suppressive assays as described in a section above.

### IL-2 expansion of EGFP^+^ Tregs in vivo

Human IL-2 (Proleukin) was diluted with NaCl for iv injection, a new dilution was used for each day of treatment, at a dose (10^4^ IU/0.62 μg/g of body weight for 4 consecutive days) previously described to expand Treg in rats (37) and analyzed at day 5 the proportion and absolute numbers of NK, CD4^+^EGFP^-^ Tconv, CD4^+^EGFP^+^ Treg, CD8^+^EGFP-Tconv and CD8^+^EGFP^+^ Treg cells in blood and spleen as well as the expression levels of Stat-5 phosphorylation (clone pY694, BD Biosciences) and CD25 (clone OX39) compared to those of these cells populations from untreated *Foxp3-EGFP* rats.

### Statistical analysis

Results are presented as means ± SD. Statistical analysis between samples was performed by a Mann-Whitney test using GraphPad Prism 4 software (GraphPad Software, San Diego, CA, USA). Differences associated with probability values of **^a^** P<0.05, **^b^** P<0.005, **^c^** P<0.0002 and **^d^**<0.0001 were considered statistically significant.

## Abbreviations

APC: Antigen Presenting Cell
APECED: autoimmune polyendocrinopathy candidiasis ectodermal dystrophy
BM: Bone Marrow
CPD: Cell Proliferation Dye
DEG: Differentially Expressed Genes
DGE: Digital Gene Expression
DNA: deoxyribonucleic acid
EDTA: Ethylenediaminetetraacetic acid
EGFP: Enhanced Green Fluorescent Protein
FDR: False Discovery Rate
FSC: forward-scattered light
GO: Gene ontologies
HCl: Hydrochloric acid
iTreg: induced regulatory T cells
iv: intravenous injection
KEGG: Kyoto Encyclopedia of Genes and Genomes
KI: Knock-In
LN: axillar Lymph Node
mAbs: monoclonal Antibody
MFI: Mean Fluorescence Intensity
MLR: Mixte Lymphocyte Reaction
NaCl: Sodium chloride
NK cells: Natural Killer cells
nTreg: natural regulatory T cells
PCA: Principal Component Analyses
PK: Proteinase K
RNA: Ribonucleic acid
SD/Crl: Sprague-Dawley
SDS: Sodium Dodecyl Sulfate
SEM: Standard Error of Mean
sgRNA: single guide RNA
SSC: side-scattered light
Tconv: Conventional T cells
TCR: T Cell Receptor
Treg: regulatory T cells
UMI: Unique Molecular Identifier
WT: Wild-Type

## Declarations

### Ethics approval and consent to participate

Not applicable.

### Consent for publication

Not applicable

### Availability of data and materials

The sequencing data have been available from the NCBI GEO database, under accession GSE189797.Graphical representations were created with R packages (35).

### Competing interests

The authors declare that they have no competing interests.

### Funding

This work was performed in the context of different programs: Biogenouest by Région Pays de la Loire, IBiSA program, TEFOR (Investissements d’Avenir French Government program, ANRII-INSB-0014), LabCom SOURIRAT project (ANR-14-LAB5-0008), Labex IGO project (Investissements d’Avenir French Government program, ANR-11-LABX-0016-01), IHU-Cesti project (Investissements d’Avenir French Government program, ANR-10-IBHU-005, Nantes Métropole and Région Pays de la Loire) and Fondation Progreffe.

### Author contributions

SM designed and performed research, analyzed data and edited the manuscript. LT performed research, analyzed data and edited the manuscript. SR, VG, CS, CU, AG, VC, L-HO, MG, J-MH and CF performed research and analyzed data. CG and JP analyzed data and edited the manuscript. IA designed research, provided funding, analyzed data and wrote the manuscript.

## Acknowledgments

This work was realized in the context of different programs: Biogenouest by Région Pays de la Loire, IBiSA program, TEFOR (Investissements d’Avenir French Government program, ANRII-INSB-0014), LabCom SOURIRAT project (ANR-14-LAB5-0008), Labex IGO project (Investissements d’Avenir French Government program, ANR-11-LABX-0016-01), IHU-Cesti project (Investissements d’Avenir French Government program, ANR-10-IBHU-005, Nantes Métropole and Région Pays de la Loire) and Fondation Progreffe. We acknowledge Bernard Malissen and Cédric Louvet for providing the *Foxp3-GFP* mice.

## Supplementary figure legends

**Supplementary figure 1. Flow cytometry analyses of spleen and thymus cells from *Foxp3-EGFP* rats. A)** Spleen and **B)** thymus were harvested from 12 weeks-old *Foxp3-EGFP* or wild-type (WT) rats and single cell suspensions were gated by SSC and FSC on lymphocytes followed by the identification with mAbs of major cell populations such as TCR+ cells, TCR+CD4+, TCR+CD8+ and CD161+ for NK cells. These populations were then analyzed for FOXP3 expression by EGFP expression and by using an anti-FOXP3 mAb. Contour plots from one animal representative of 6 analyzed in the same conditions.

**Supplementary figure 2. Suppression assay using T CD4^+^EGFP^+^ and CD8^+^EGFP^+^ cells.** Representative histograms of a suppressive assay using spleen cells from 12 weeks-old *Foxp3-EGFP* rats. TCR^+^CD4^+^CD25^+^CD127^low^EGFP^+^ (labeled with CPD-450) and TCR^+^CD8^+^CD45RC^low/-^EGFP^+^ cells were sorted and added to an MLR in a 1:1 or 1:2 Tregs to Tconv cells. The MLR was performed by co-culturing in a 1:1 ratio spleen CD4^+^CD25^-^ Tconv cells from SPD rats (MHC haplotype u) labelled with CPD-670 along with enriched spleen APCs from Lewis 1A (MHC haplotype a) rats. Decrease in the percentage of CPD-670 bright cells and increase in the percentages of CPD-670 low cells denotes dilution of CPD-670 with cell proliferation. The left histogram shows the CPD-670 dilution of proliferating T cells in the absence of Treg and the others show less proliferation in the presence of the EGFP^+^ Treg. Representative experiment out of 4 performed.

**Supplementary figure 3. RNAseq analyses of EGFP^+^ and EGFP^-^ within CD4^+^ and CD8^+^ T cells.** TCR^+^CD25^high^CD127^low^CD4^+^ and TCR^+^CD45RC^low/-^ CD8^+^ EGFP^+^ and EGFP^-^ T cells were cell sorted and RNAseq analyses were performed comparing CD4^+^EGFP^+^ vs. CD4^+^EGFP^-^ cells, and CD8^+^EGFP^+^ vs. CD8^+^EGFP^-^ cells and CD8^+^EGFP^+^ vs. CD4^+^EGFP^+^ cells. **A.** R-Shiny representations of RNAseq levels for individual samples of the indicated genes with immune functions expressed in units per million reads. **B.** GO analyses of analysis of functional pathways, Cell component analysis showed that the proteins encoded by comparison of TCR^+^CD25^high^CD127^low^CD4^+^EGFP^+^ vs. TCR^+^CD25^high^CD127^low^CD4^+^EGFP^-^, Comparison of 5 principal pathways in immune cell activation **(b)** Comparison of 3 principal pathway in immune cell proliferation. **(c)** Comparison of 4 principal pathway in immune cell adhesion. **(d)** Comparison of 11 pathways in miscellaneous process. **C.** Venn diagrams showing the overlap in genes that were significantly (adjusted p value, 0.05 and absolute log2 fold-change of genes differentially upregulated (left) or down regulated (right).

**Supplementary figure 4. Confirmation of some RNAseq results on CD4+EGFP+ and CD8+EGFP+ cells by FACS. A.** Histograms of TCR+CD4+EGFP+, TCR+CD4+EGFP-, TCR+CD8+EGFP+and TCR+CD8+EGFP-cells analyzed with the indicate mAbs. Percentages above the traits correspond to the number of positive cells above staining with isotype control mAbs. One experiment representative of 3 performed in the same conditions. **B.** R-Shiny representation of RNAseq expression levels expressed in individual sample in units per million reads for *Il2ra (Cd25), Cd44 and Icos*.

**Supplementary figure 5. IL-2 induce expansion of EGFP^+^ Tregs in spleen.** *Foxp3-EGFP* rats received or not hIL-2 at low doses and the percent of the indicated cell populations was analyzed in spleen. Percentages of cells in the indicated cell populations of IL-2-treated and untreated rats. Left graph shows mean +/− SEM, n=3, and right contour plots show a representative animal.

**Supplementary table 1.** Frequency of CD4^+^EGFP^+^ and CD8^+^EGFP^+^ in *Foxp3-EGFP* rats and *Foxp3-EGFP* mice.

|  |  | thymus | LN | spleen | BM | blood/ml |
| --- | --- | --- | --- | --- | --- | --- |
| **Rat\*\*** | **TCR<sup>+</sup>CD4<sup>+</sup>EGFP<sup>+</sup>** | 5.2 ± 0.9** | 7.8 ± 1.7 | 10 ± 0.2 | 7.2 ± 3 | 7.9 ± 0.6 |
|  | **TCR<sup>+</sup>CD8<sup>+</sup>EGFP<sup>+</sup>** | 0.6 ± 0.15 | 1.3 ± 0.4 | 0.7 ± 0.01 | 0.8 ± 0.45 | 0.7 ± 0.1 |
| **Mouse\*\*** | **TCR<sup>+</sup>CD4<sup>+</sup>EGFP<sup>+</sup>** | 0.18 ± 0.06 | 3.9 ± 3.3 | 9.7 ± 2.5 | 7.9 ± 4.1 | 3.7 ± 1.0 |
|  | **TCR<sup>+</sup>CD8<sup>+</sup>EGFP<sup>+</sup>** | 0.003 ± 0.005 | 0.3 ± 0.6 | 0.3 ± 0.2 | 0.06± 0.09 | 0.4 ± 0.3 |

**\*N=3 for rats and n=5 for mice**
**\*\*% of total cells**

**Supplementary table 2.**
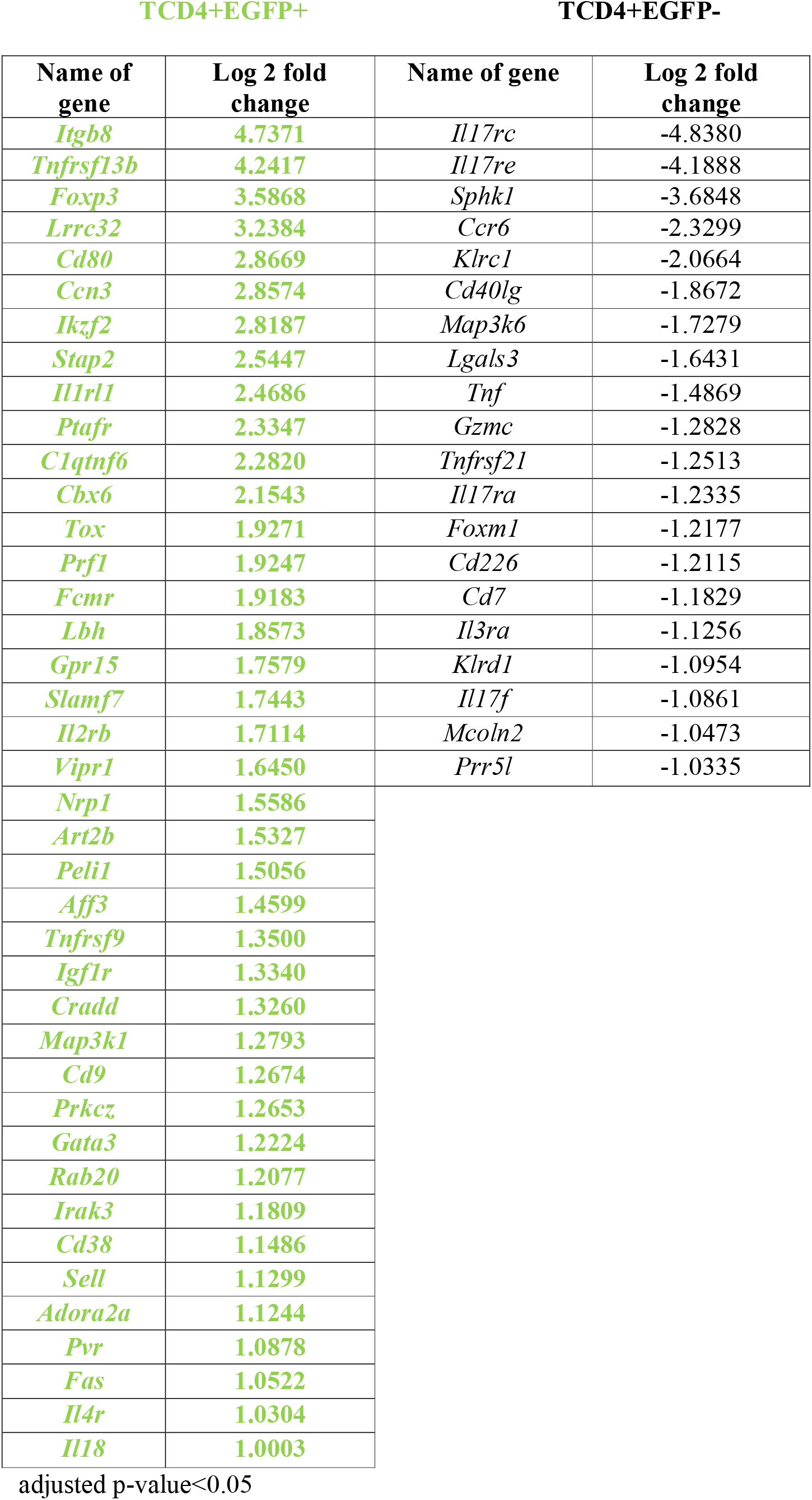
Immune genes upregulated in TCD4+EGFP+ vs. TCD4+EGFP-cells.

**Supplementary table 3.**
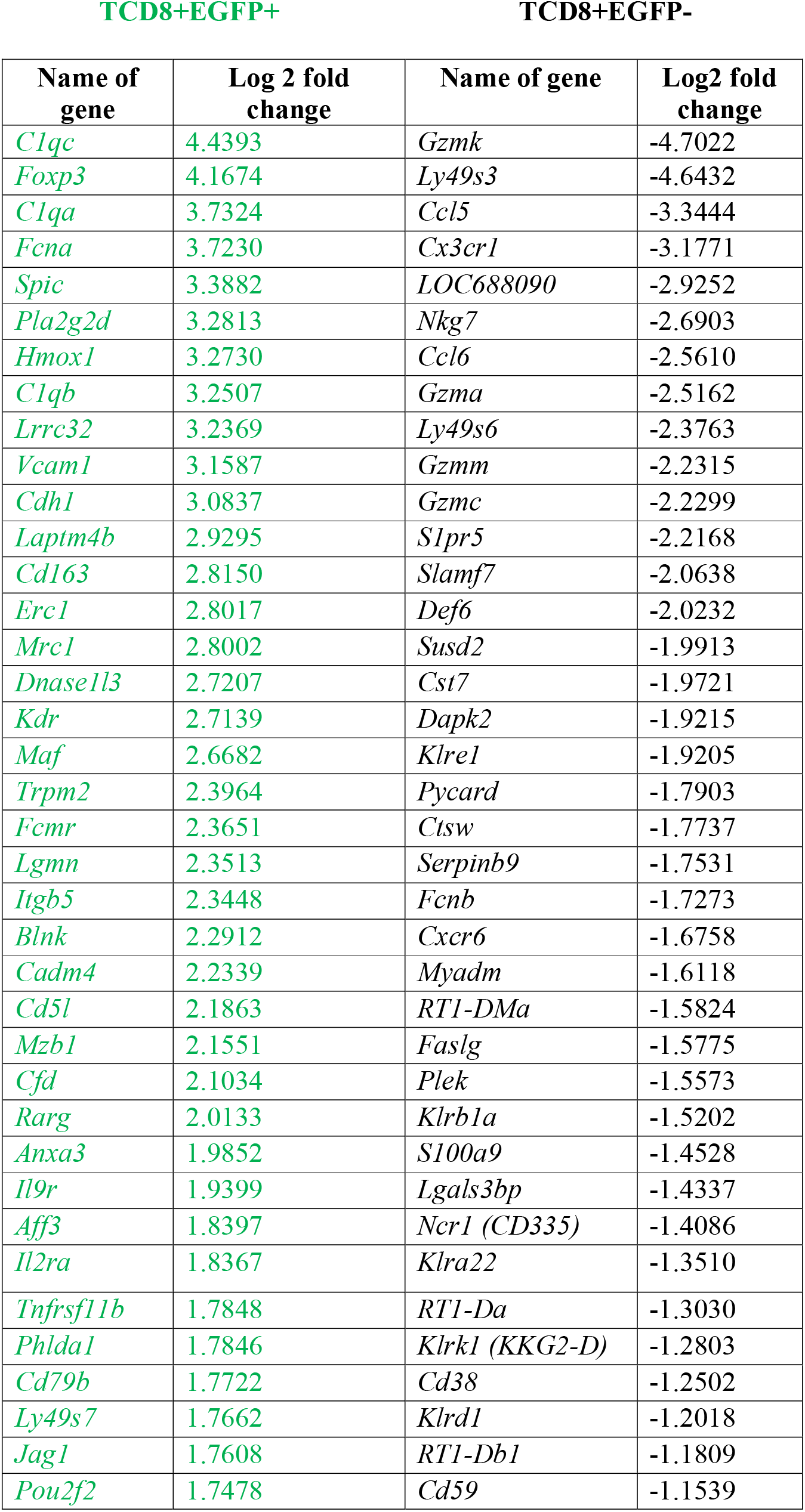
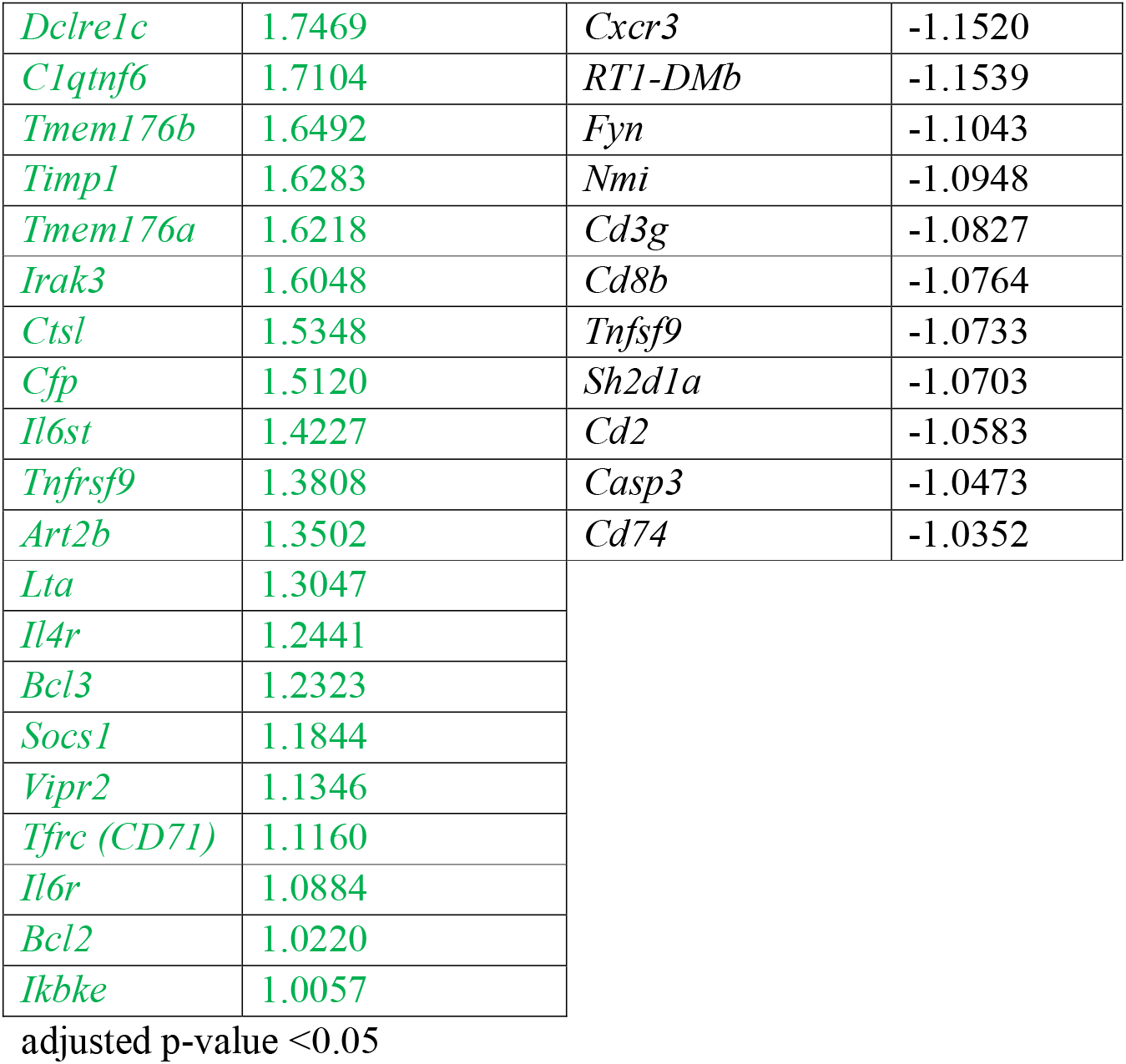
Immune genes upregulated in TCD8+EGFP+ vs. TCD8+EGFP-cells.

**Supplementary table 4.**
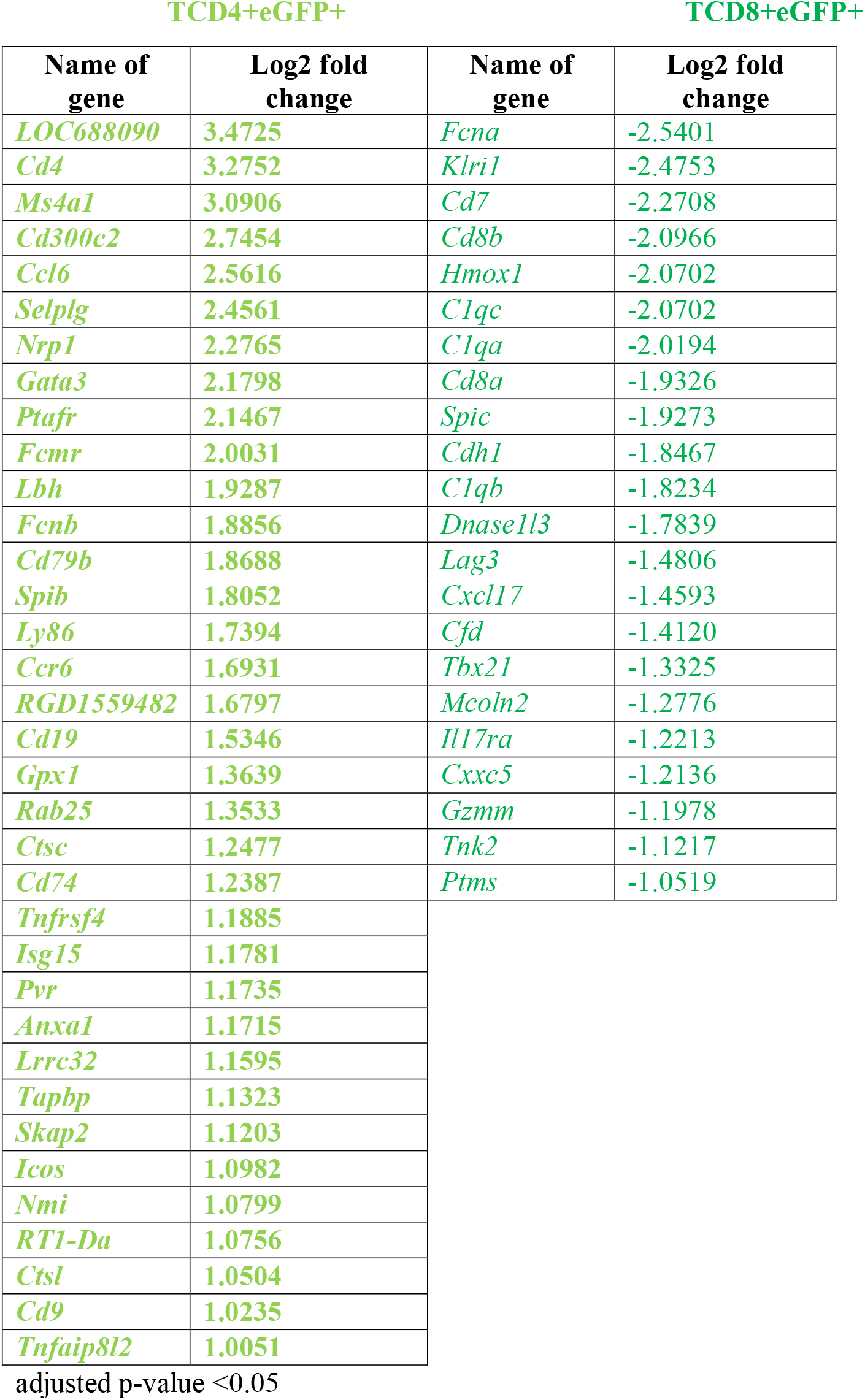
Immune genes upregulated in CD4+EGFP+ vs. CD8+EGFP+ cells.

**Supplementary table 5.**
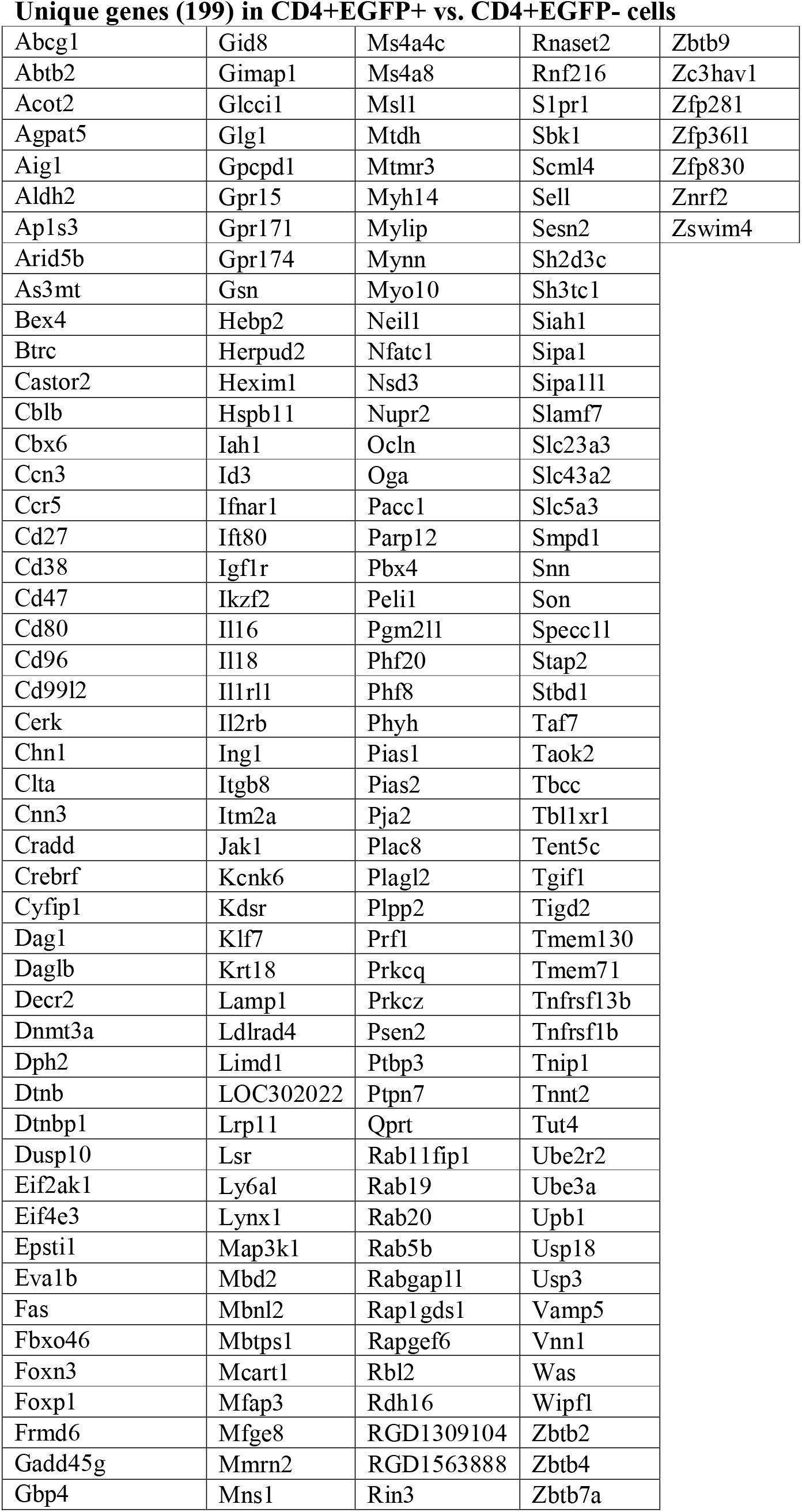
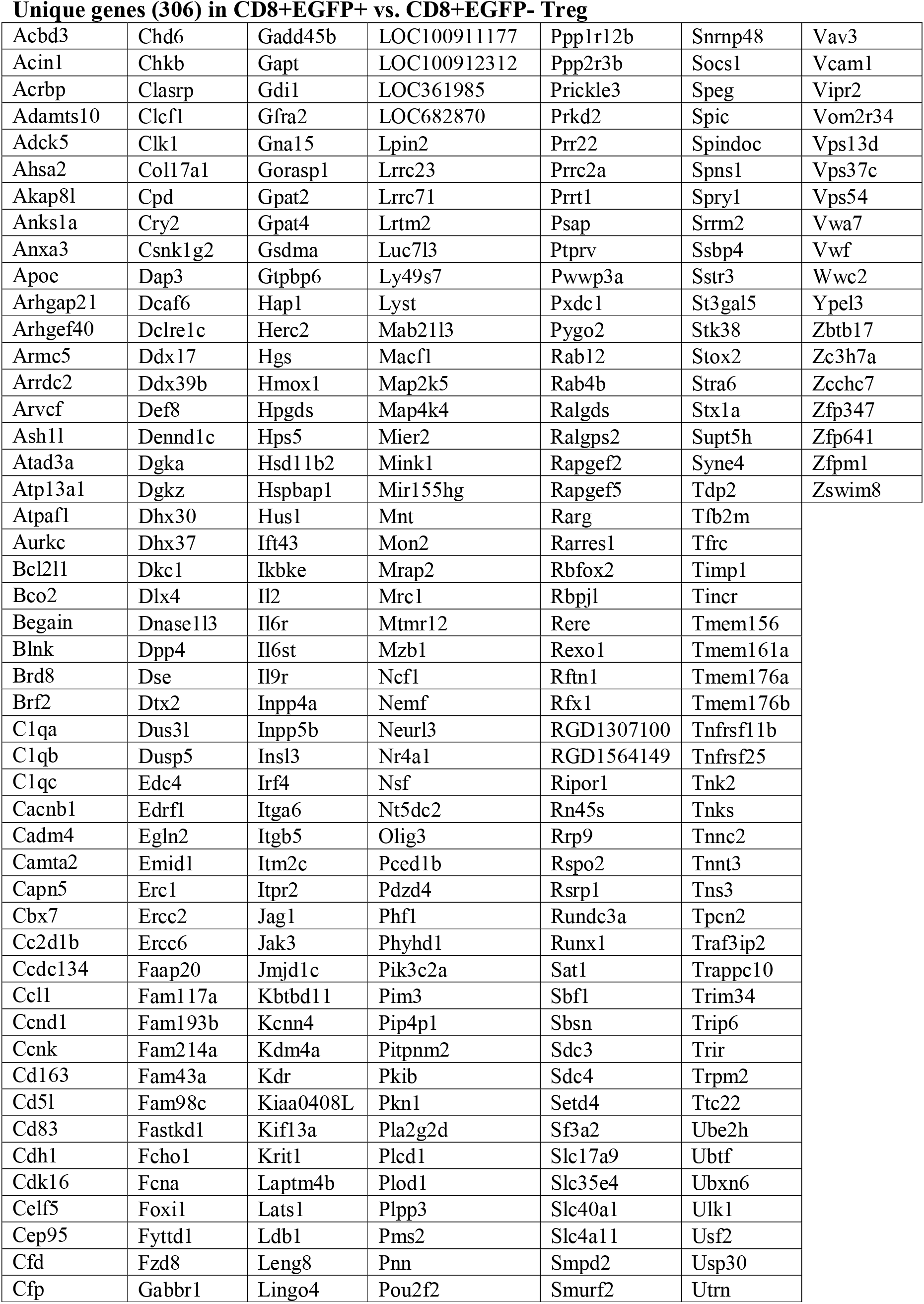
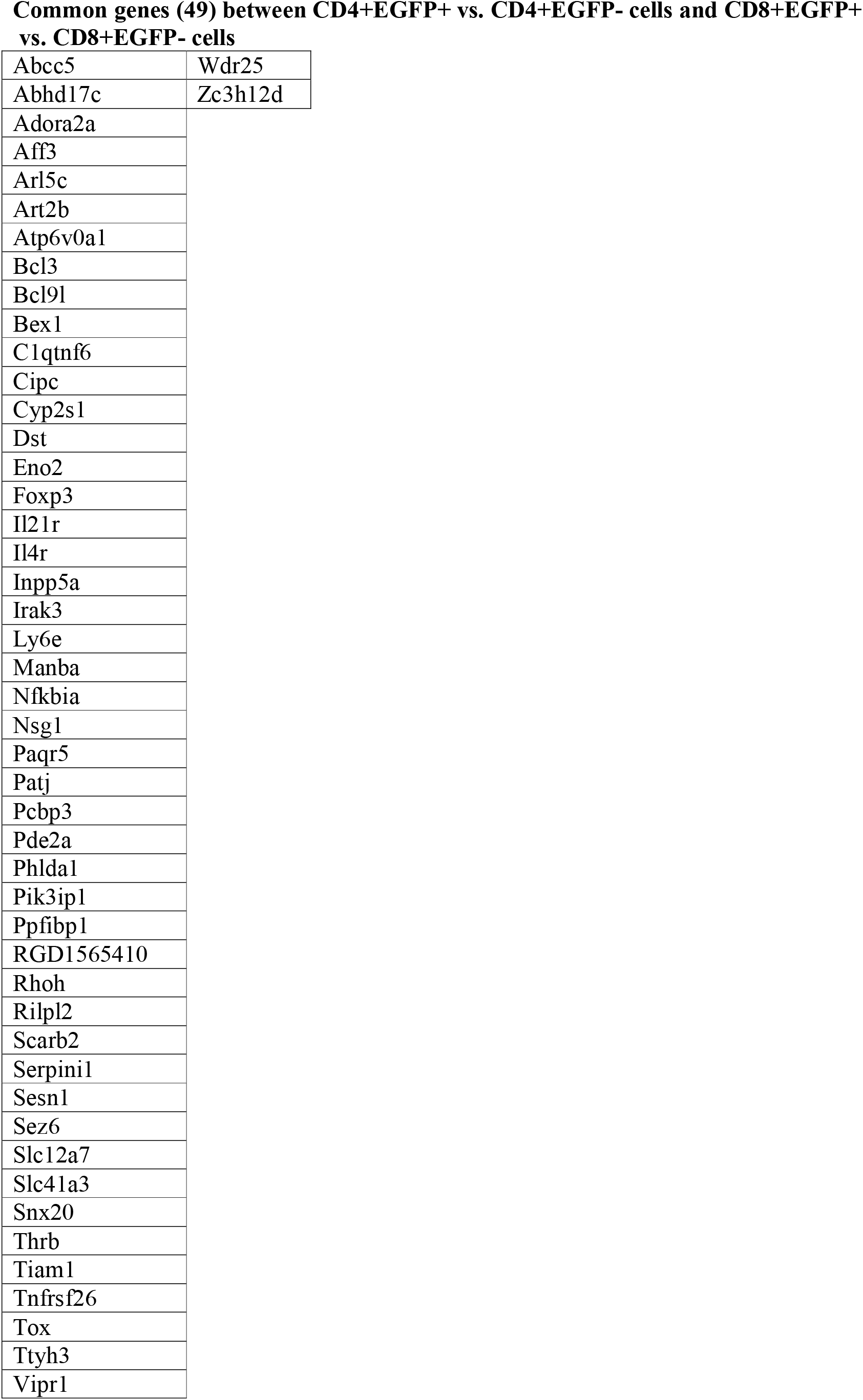
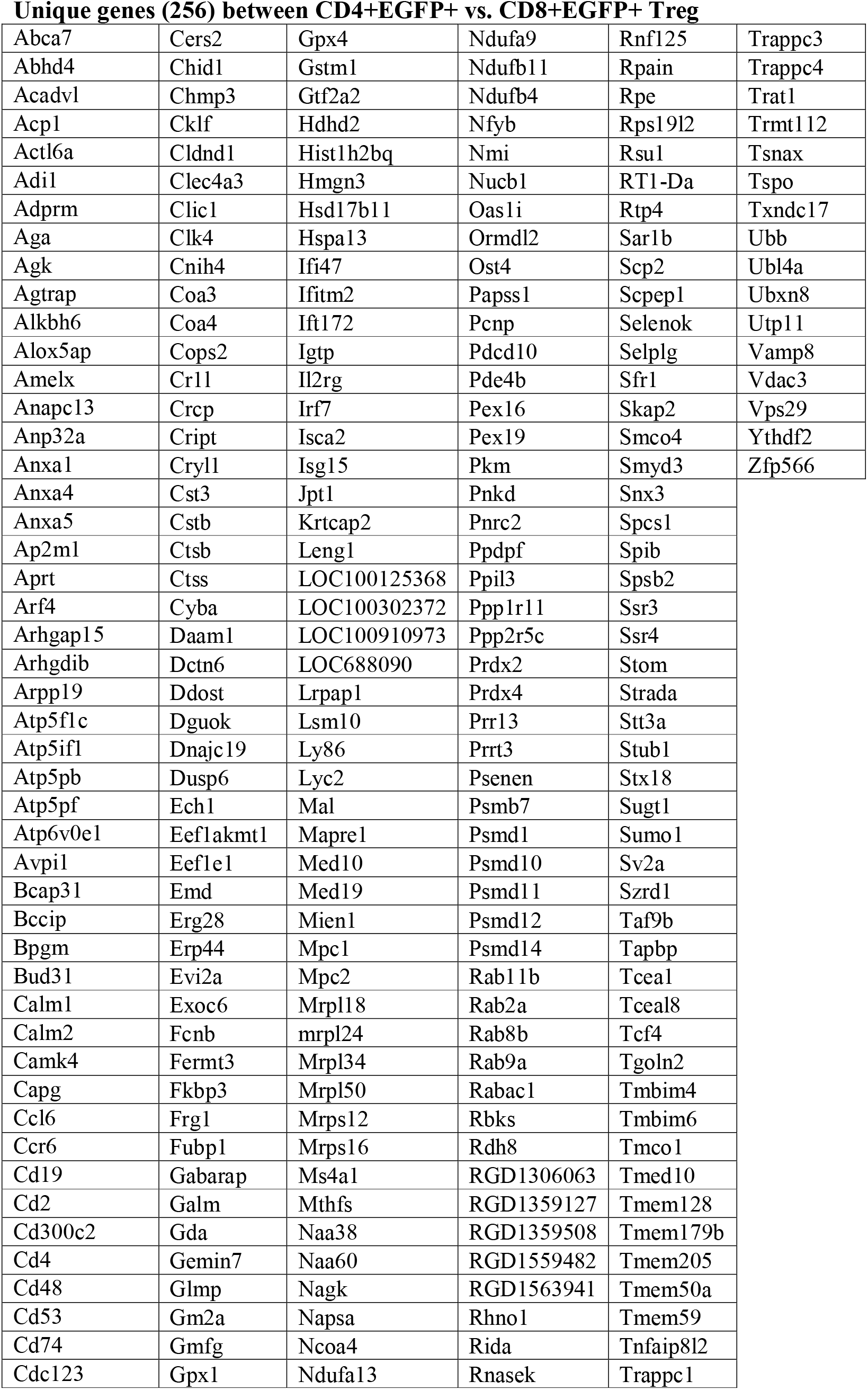
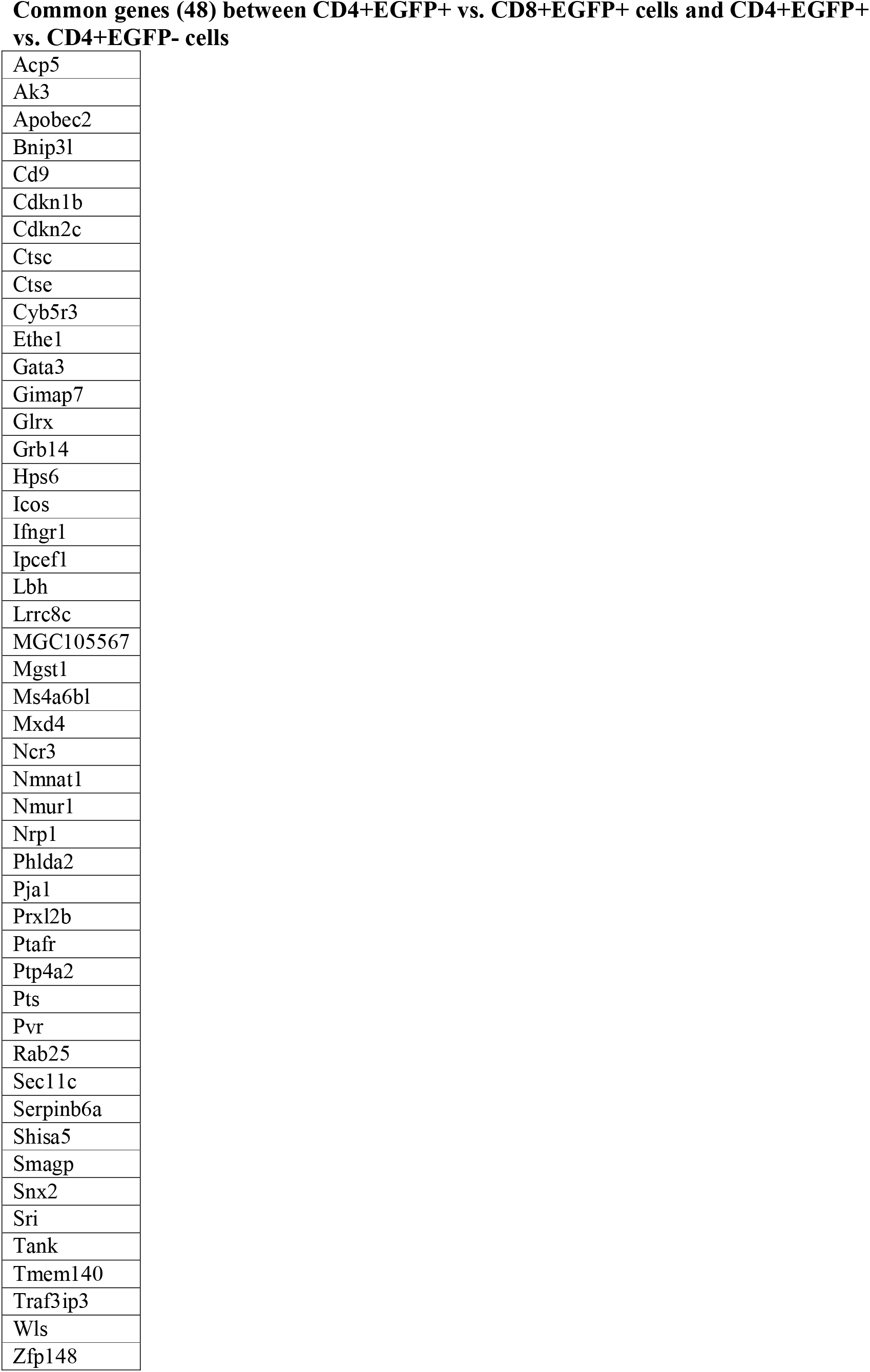
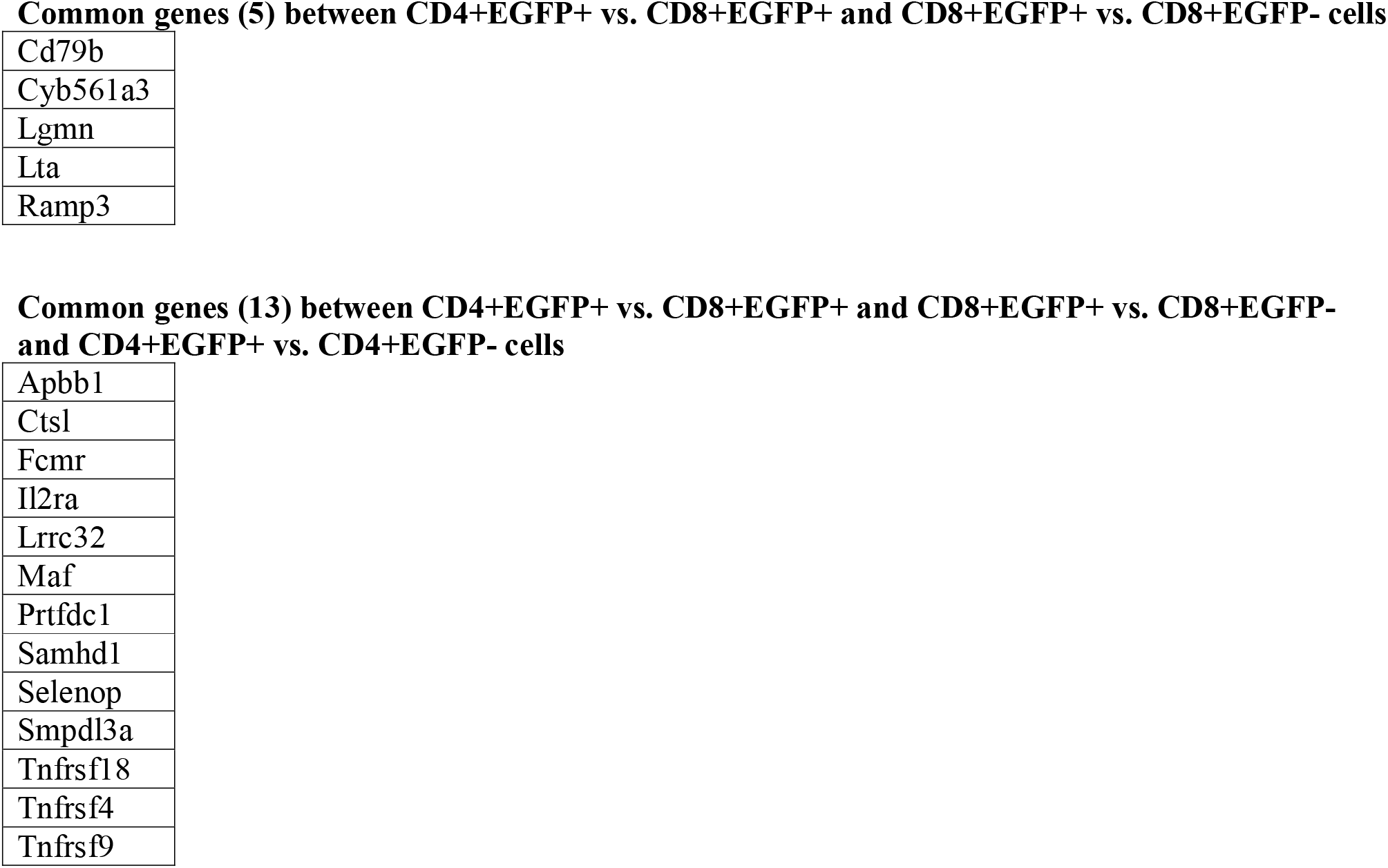
Genes that are upregulated in Venn diagrams.

**Supplementary table 6.**
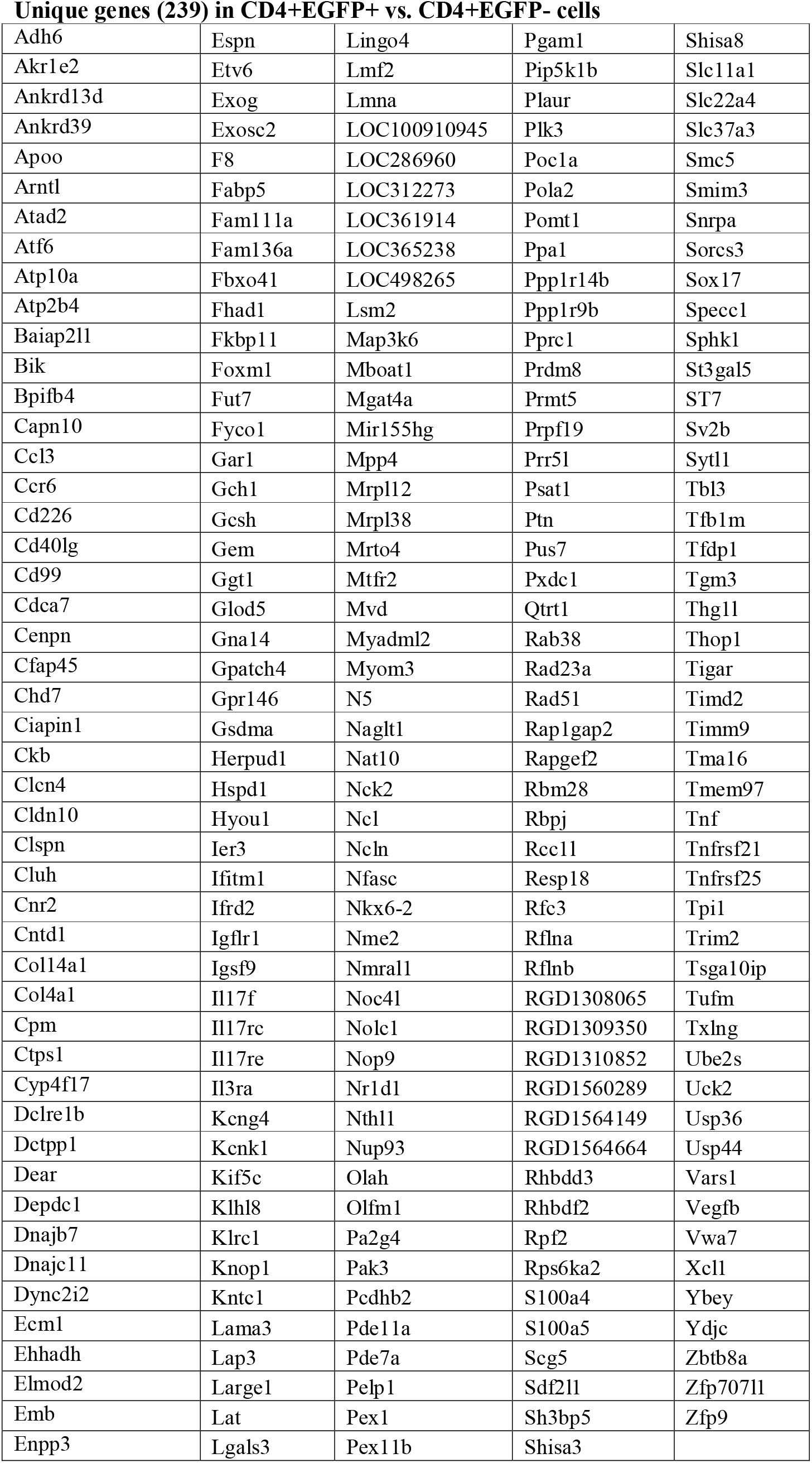
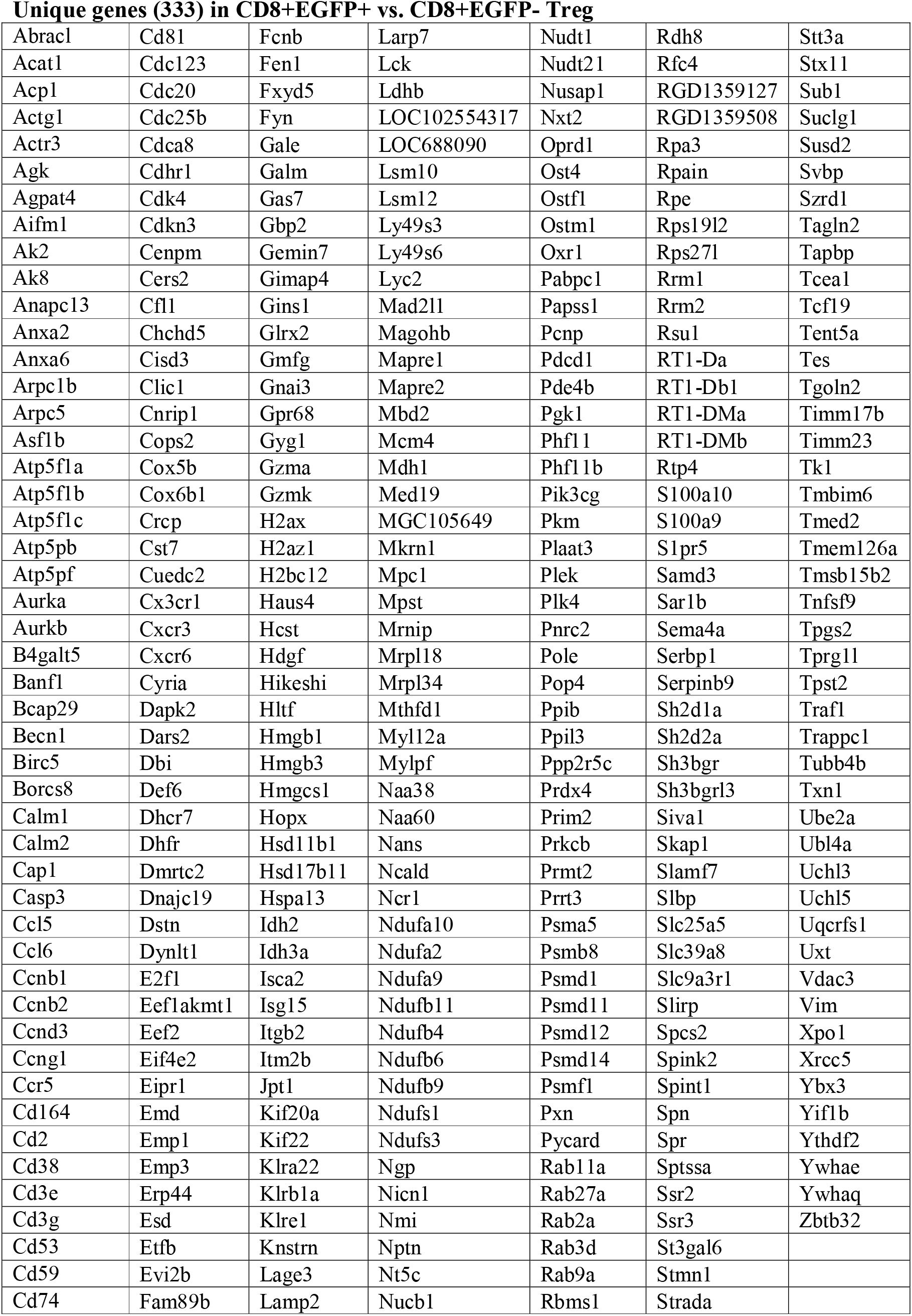
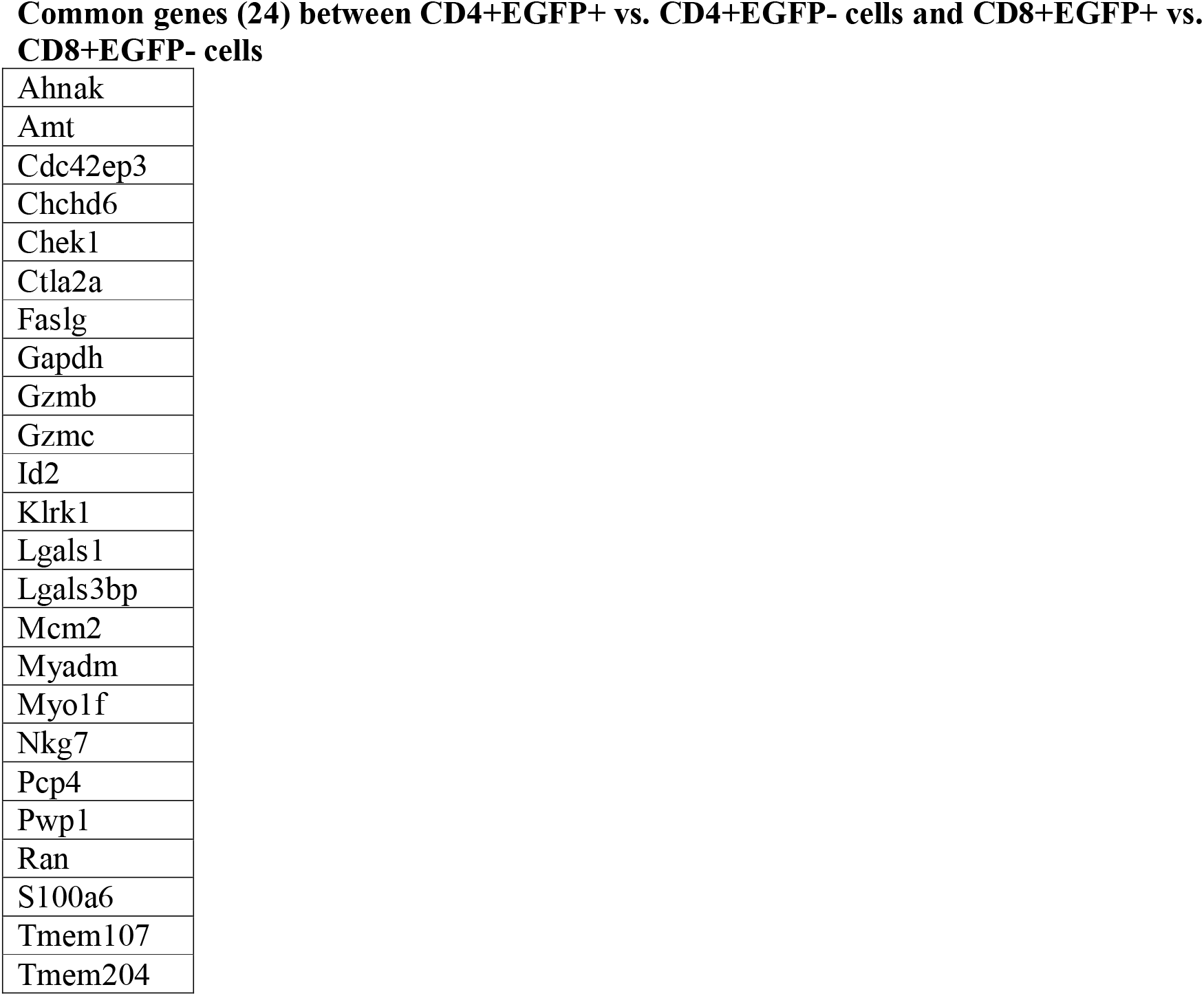
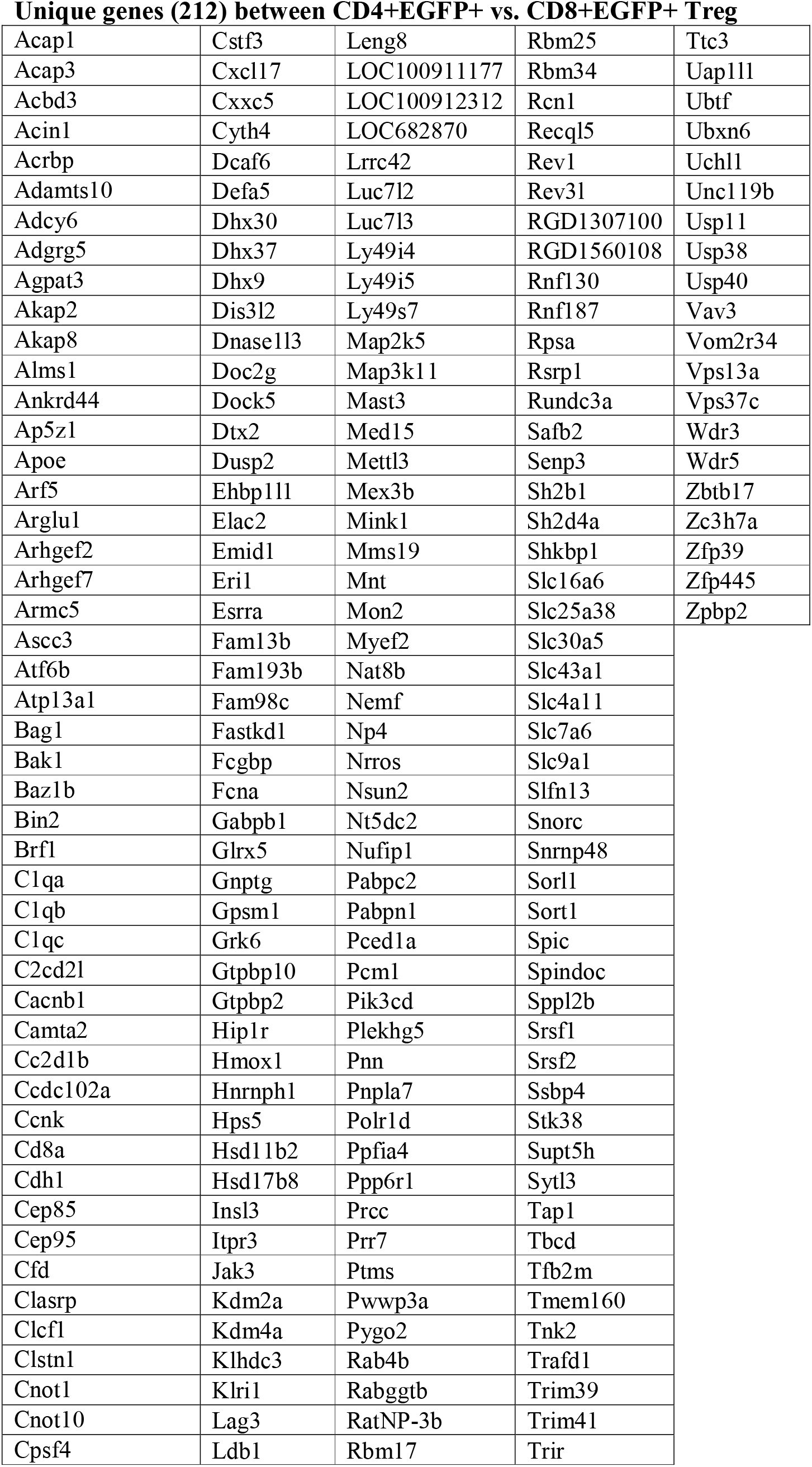
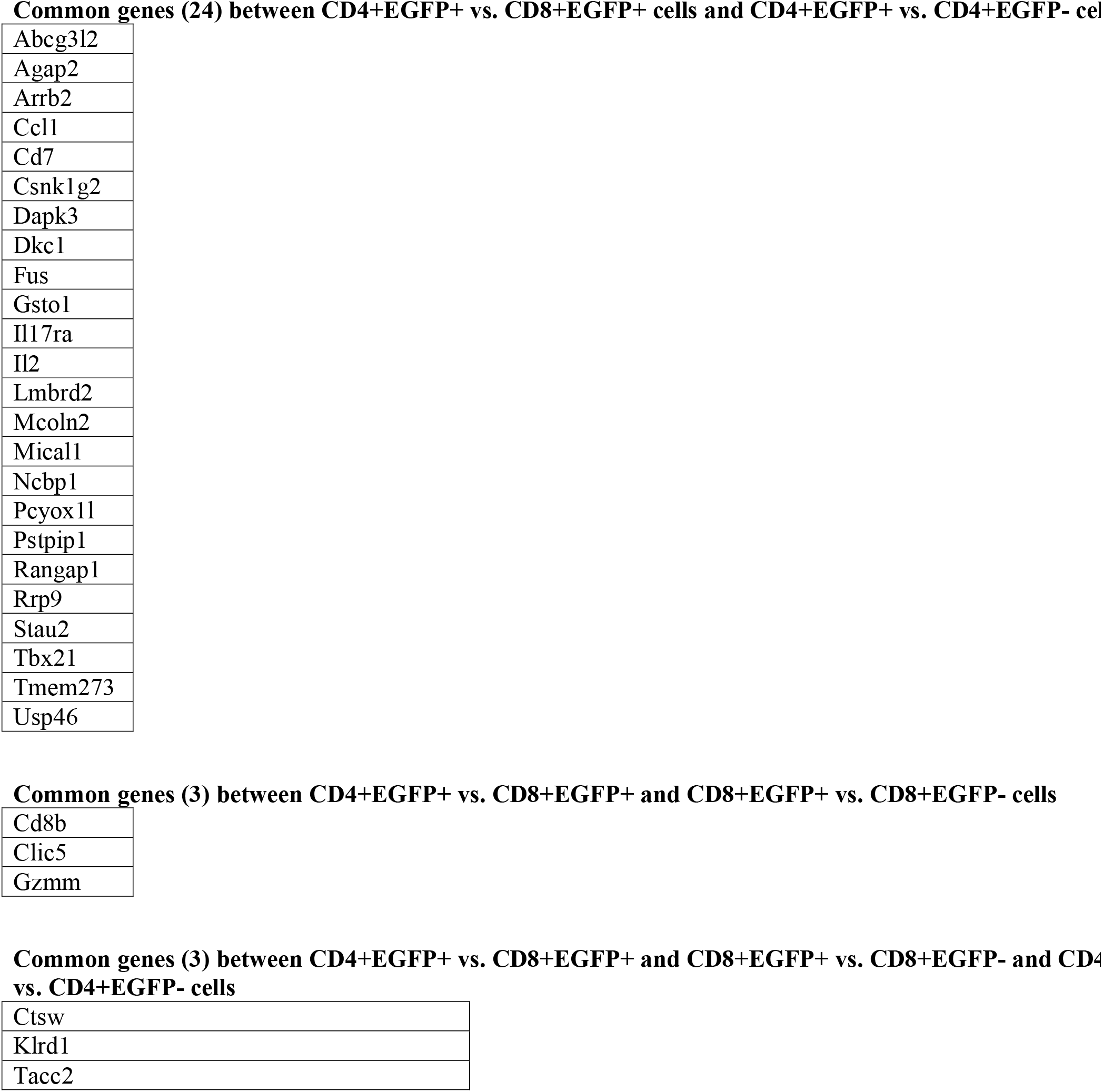
Genes that are downregulated in Venn diagrams.

**Supplementary table 7.** Increase in different EGFP+ and EGFP-cell subsets in blood following IL-2 treatment.

|  |  |  | **CD3+ CD4+**<br>**EGFP+** | **CD3+CD4+**<br>**EGFP-** | **CD3+CD8+**<br>**EGFP+** | **CD3+CD8+**<br>**EGFP-** | **CD161+**<br>**EGFP+** | **CD161+**<br>**EGFP-** |
| --- | --- | --- | --- | --- | --- | --- | --- | --- |
| **Blood** | **Untreated**<br>**n=4** | **% of total**<br>**cells** | 5.6 ± 1.4 | 94.9 ± 0.3 | 2.2 ± 0.4 | 96 ± 0.3 | 0 | 2 ± 1.4 |
|  |  | **N°cellsx10<sup>5</sup>**<br>**/ml** | 0.08 ± 0.06 | 1.7 ± 0.6 | 0.016 ±<br>0.005 | 0.7 ± 0.3 | 0 | 0.06 ±<br>0.04 |
|  | **hIL2**<br>**treated**<br>**n=3** | **% of total**<br>**cells** | 17.1 ± 0.6 | 82.6 ± 0.4 | 9.8 ± 0.9 | 89.7 ± 0.9 | 0 | 2,7 ± 1.1 |
|  |  | **N°cellsx10<sup>5</sup>**<br>**/ml** | 1.5 ± 0.5 | 7.5 ± 2 | 0.5 ± 0.1 | 4.4 ± 1.5 | 0 | 0.6 ± 0.2 |
|  |  | **Fold**<br>**induction\*** | 19 | 4.4 | 31 | 6.3 | 0 | 10 |

\* Fold induction of absolute numbers of cells in the IL-2-treated vs. untreated animals

**Supplementary table 8.** Antibodies used in the study.

| **Marker** | **Clone** | **Provider** |
| --- | --- | --- |
| Viability Dye 506 |  | Invitrogen eBiosciences |
| Biotin-conjugated mouse anti-rat CD3 | G4.18 | BD Biosciences |
| BV421-conjugated mouse anti-rat CD3 | 1F4 | BD Biosciences |
| PercP-conjugated mouse anti-rat TCR$\alpha\beta$ | R7.3 | BD Biosciences |
| PECy7-conjugated mouse anti-rat CD4 | OX35 | BD Biosciences |
| BV421-conjugated mouse anti-rat CD25 | OX39 | BD Biosciences |
| APC-conjugated mouse anti-rat CD161 | 3.2.3 | Produced in house |
| APC-conjugated mouse anti-rat CD8a | OX8 | Produced in house |
| PE-conjugated mouse anti rat CD45RC | OX22 | BD Biosciences |
| PE-conjugated anti-Foxp3 | FJK-16s | Invitrogen eBiosciences |
| PE-conjugated mouse anti-Stat5 | pY694 | BD Biosciences |
| PE-conjugated mouse anti-rat RT1B | OX6 | BD Biosciences |
| PE-conjugated mouse anti-rat CD28 | CD28 | BD Pharmingen |
| PE-conjugated hamster anti-mouse CD27 | LG.3A10 | BD Biosciences |
| BV711-conjugated mouse anti-rat CD44 | OX49 | BD Biosciences |
| BV421-conjugated hamster anti-rat CD62L | HRL1 | BD Biosciences |
| Purified mouse anti-rat CD71 | OX26 | Produced in house |
| Biotin-conjugated mouse anti-rat CD5 | OX19 | Produced in house |
| Purified mouse anti-rat CD26 | OX61 | Produced in house |
| Purified rabbit anti- rat CX3CR1 | TP501 | Produced in house |
| PE-conjugated mouse anti-rat CD106 | MR106 | BD Biosciences |
| PE-conjugated mouse anti-rat CD80 | 3H5 | BD Biosciences |
| Purified mouse anti-rat CD38 | 14.27 | Biolegend |
| PE-conjugated or BV421-conjugated mouse anti-human CD137 | 4BB1 | BD Biosciences |
| Streptavidin APC-Cy<sup>TM</sup>7 |  | BD Biosciences |
| AF568 goat anti-mouse IgG2a |  | Thermofisher |
| R-Phycoerythrin AffiniPure F(ab')<sub>2</sub> □<br>Fragment Donkey Anti-Rabbit IgG (H+L) |  | Jackson Immunoresearch |

## Notes

### Competing Interest Statement

The authors have declared no competing interest.

