## Supplemental figures for "CD4^+^ and CD8^+^ regulatory T cells characterization in the rat using a unique transgenic *Foxp3-EGFP* model"

#### Supplementary figure 1.

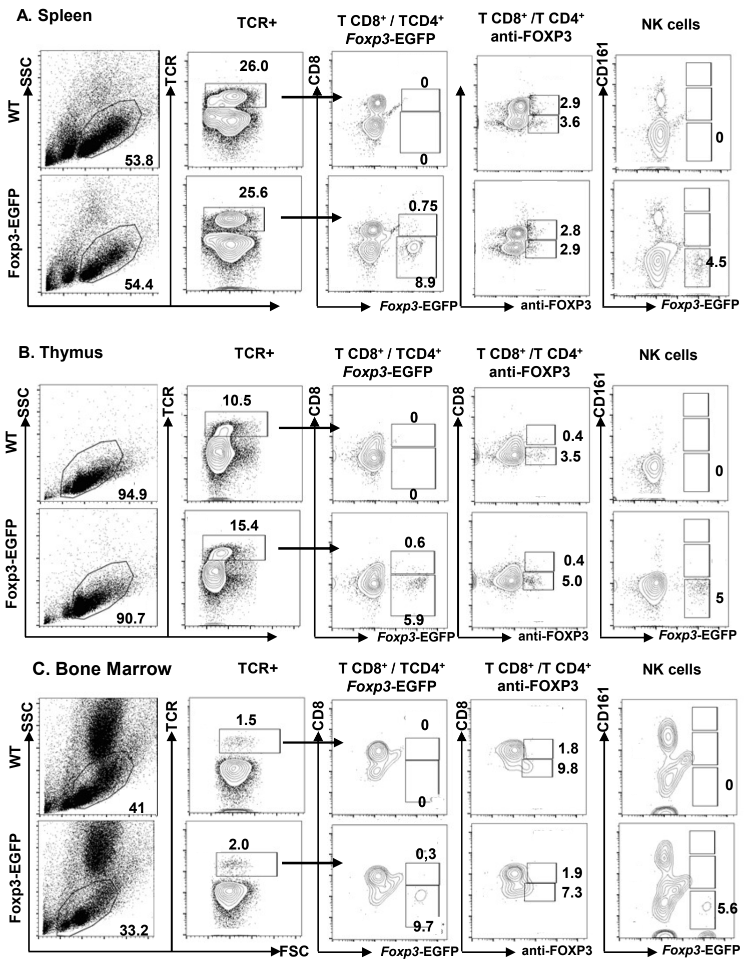

### Supplementary figure2.

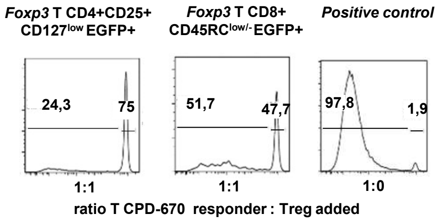

#### Supplementary figure 3.

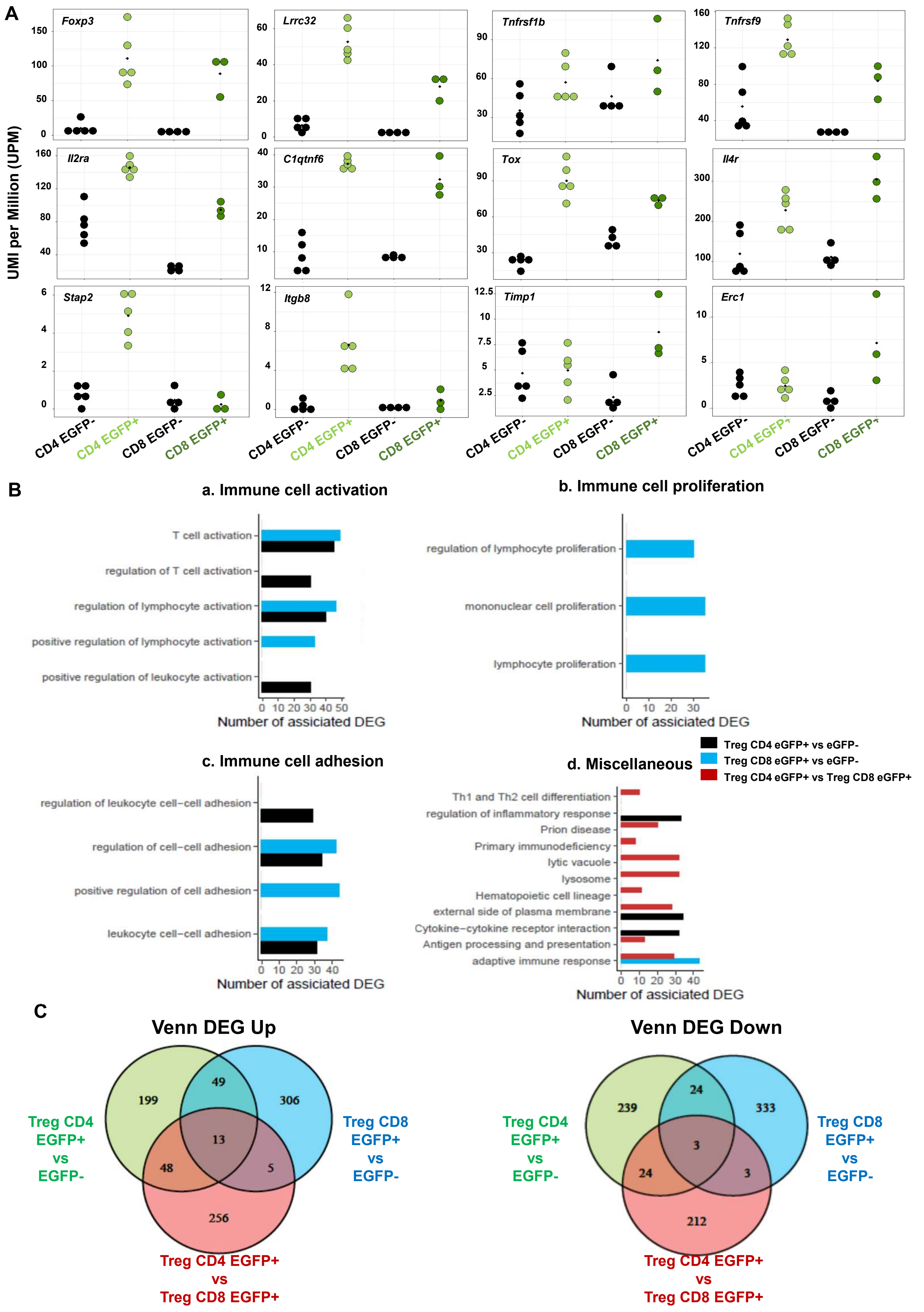

**Supplementary figure 4. Confirmation of RNAseq results at the protein level by cytometry.**

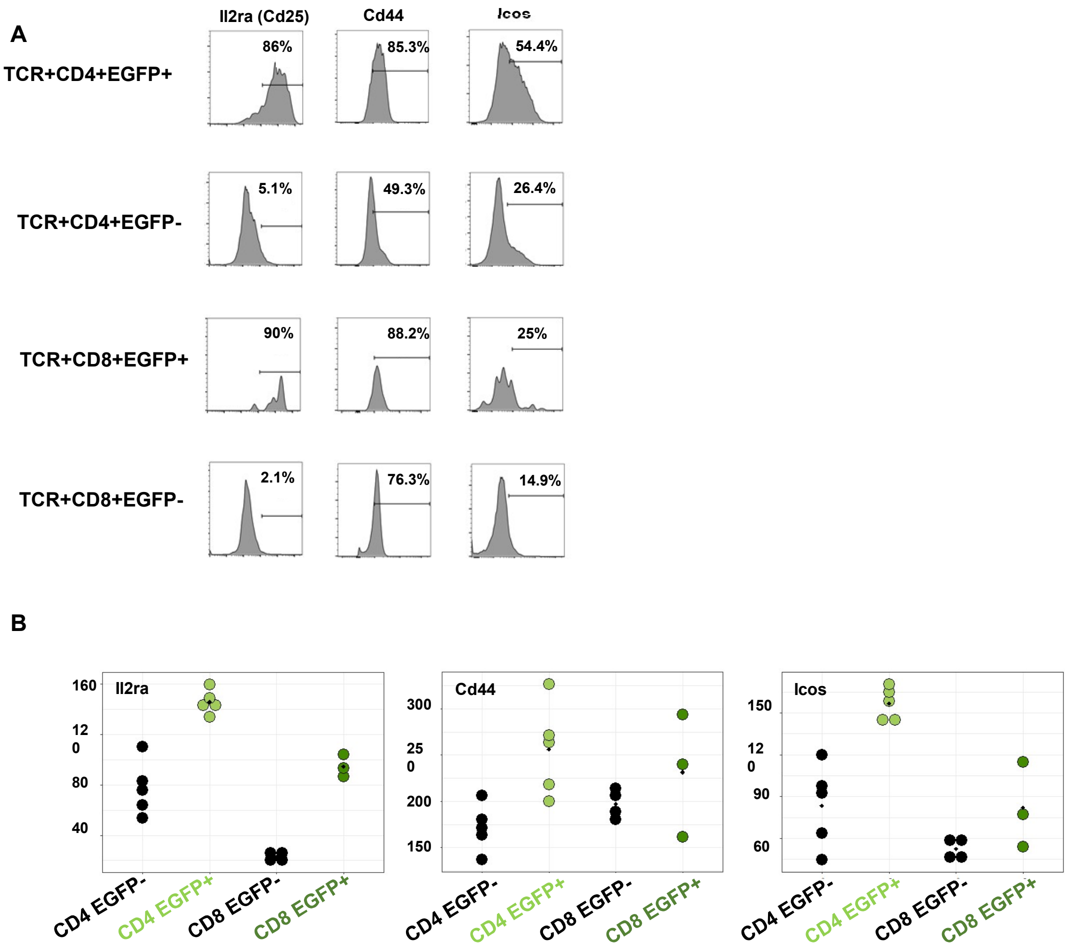

#### Supplementary figure 5.

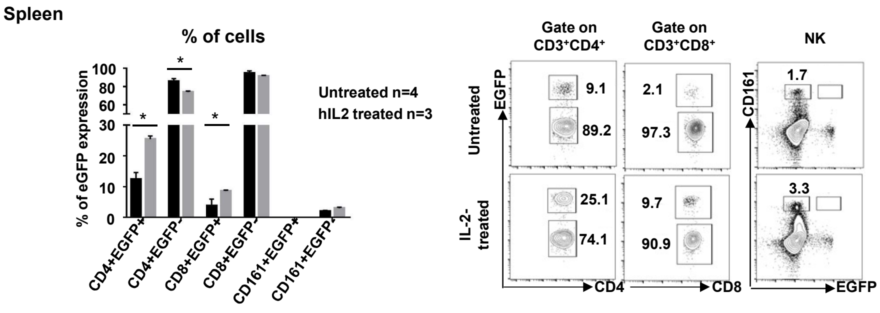
